# A genetic screen for dominant chloroplast reactive oxygen species signaling mutants reveals life stage-specific singlet oxygen signaling networks

**DOI:** 10.1101/2023.10.26.564295

**Authors:** Matthew D. Lemke, Jesse D. Woodson

## Abstract

Plants employ intricate molecular mechanisms to respond to abiotic stresses, which often lead to the accumulation of reactive oxygen species (ROS) within organelles such as chloroplasts. Such ROS can produce stress signals that regulate cellular response mechanisms. One ROS, singlet oxygen (^1^O_2_), is predominantly produced in the chloroplast during photosynthesis and can trigger chloroplast degradation, programmed cell death (PCD), and retrograde (organelle-to-nucleus) signaling. However, little is known about the molecular mechanisms involved in these signaling pathways or how many different signaling ^1^O_2_ pathways may exist. The *Arabidopsis thaliana plastid ferrochelatase two* (*fc2*) mutant conditionally accumulates chloroplast ^1^O_2_, making *fc2* a valuable genetic system for studying chloroplast ^1^O_2_-initiated signaling. Here, we have used activation tagging in a new forward genetic screen to identify eight dominant *fc2* activation-tagged (*fas*) mutations that suppress chloroplast ^1^O_2_-initiated PCD. While *fc2 fas* mutants all block ^1^O_2_-triggered PCD in the adult stage, only two *fc2 fas* mutants block such cellular degradation at the seedling stage, suggesting that life-stage-specific ^1^O_2_-response pathways exist. In addition to PCD, *fas* mutations generally reduce ^1^O_2_-induced retrograde signals. Furthermore, *fas* mutants have enhanced tolerance to excess light, a natural mechanism to produce chloroplast ^1^O_2_. However, general abiotic stress tolerance was only observed in one *fc2 fas* mutant (*fc2 fas2*). Together, this suggests that plants can employ general stress tolerance mechanisms to overcome ^1^O_2_ production but that this screen was mostly specific to ^1^O_2_ signaling. We also observed that salicylic acid (SA) and jasmonate (JA) stress hormone response marker genes were induced in ^1^O_2_-stressed *fc2* and generally reduced by *fas* mutations, suggesting that SA and JA signaling is correlated with active ^1^O_2_ signaling and PCD. Together, this work highlights the complexity of ^1^O_2_ signaling by demonstrating that multiple pathways may exist and introduces a suite of new ^1^O_2_ signaling mutants to investigate the mechanisms controlling chloroplast-initiated degradation, PCD, and retrograde signaling.

## Introduction

Abiotic environmental stresses can severely and negatively affect plant fitness and, consequently, agricultural yields (Kopecka et al., 2023). As sessile organisms, plants have evolved elaborate signaling mechanisms that allow them to sense environmental changes and acclimate. For instance, plant cells can use their energy-producing organelles (chloroplast and mitochondria) for such purposes. Under stress, the chloroplast can produce retrograde (chloroplast-to-nucleus) signals to regulate nuclear-encoded genes, including those involved in photosynthesis and acclimation (Dogra and Kim, 2019, Chan et al., 2016, De Souza et al., 2017). Although the mechanisms behind these signals are poorly understood, some have been shown to involve the production of reactive oxygen species (ROS) within chloroplasts following abiotic stress. Plants have sophisticated systems to mediate such ROS accumulation, including enzymatic and chemical quenching via ROS scavengers (You and Chan, 2015) and pigments (Foyer, 2018), respectively. When these systems become overwhelmed, ROS can damage cellular components, induce photo-inhibition, and lead to cellular degradation (Foyer, 2018). In the latter case, it is becoming clear that cellular degradation can be genetically initiated by ROS and lead to programmed cell death (PCD) and chloroplast quality control (CQC). PCD pathways can limit systematic damage to local tissue and help prevent water loss, CQC pathways help maintain healthy populations of photosynthesizing chloroplasts, and both pathways help to remobilize nutrients to healthy plant tissue (van Doorn and Woltering, 2004, Cruz de Carvalho, 2008, Woodson, 2022). ROS are naturally produced during photosynthesis when excess light (EL) energy is absorbed by chlorophyll (Triantaphylidès et al., 2008). Plants manage excess light energy through photochemical reactions (photosynthesis), the release of photons via chlorophyll fluorescence, or heat dissipation via nonphotochemical chlorophyll fluorescence quenching (NPQ) (Ruban, 2016).

Excess energy not dissipated by these mechanisms can lead to the formation of ROS. This includes singlet oxygen (^1^O_2_), produced at photosystem II (PSII), and superoxide (O_2_^-^) and hydrogen peroxide (H_2_O_2_), produced at photosystem I (PSI) (Triantaphylidès and Havaux, 2009, Krieger-Liszkay et al., 2011). Specifically, ^1^O_2_ is generated by the transfer of energy from excited chlorophylls to ground-state oxygen (Triantaphylidès and Havaux, 2009). ^1^O_2_ is highly reactive, has a short half-life (< 200 ns) (Gorman and Rodgers, 1992), and has an expected diffusion distance of < 155 nm (Ogilby, 2010). Thus, the bulk of ^1^O_2_ generated within a chloroplast (an organelle that is 2-3 μm wide and 5-10 μm long) is likely to remain compartmentalized to this organelle and oxidize chloroplast macromolecules, including the proteinaceous photosynthetic machinery, lipids, nucleic acids, and carotenoids, leading to chloroplast dysfunction.

While very high levels of ^1^O_2_ are toxic to the cell, studies using *Arabidopsis thaliana* have demonstrated that cellular degradation can be a genetically controlled response to ^1^O_2_ (Wagner et al., 2004, Ramel et al., 2013, Woodson et al., 2015). The Arabidopsis *fluorescent in blue light* (*flu*) mutant is defective in a gene encoding a chloroplast-membrane-bound protein that negatively regulates the Mg^+2^ branch of the tetrapyrrole pathway and accumulates protochlorophyllide (Pchlide), a chlorophyll precursor, in the dark (Meskauskiene et al., 2001). Upon a dark-to-light shift, these mutants bleach and die due to the photo-excitation of excess Pchlide, which produces a burst of ^1^O_2_ within chloroplasts. This ^1^O_2_ induces ^1^O_2_ response genes (SORGs), followed by programmed cell death, a response that was shown to be dependent on a chloroplast retrograde signal involving the Executor 1 (EX1) protein (Wagner et al., 2004).

Another mutant, *chlorina* (*ch1*), lacks chlorophyll *a* oxygenase activity, thereby reducing the production of chlorophyll *b*. Consequently, these mutants have impaired photoprotection and generation of excess ^1^O_2_ at the PSII reaction centers (Ramel et al., 2013). This ^1^O_2_ production leads to sensitivity to EL, triggering PCD and retrograde signaling. This PCD is also regulated by a genetic signal, as evidenced by two mutations, *plant u-box 4* (*pub4-6*) and *oxidative signal inducible 1*(*oxi1*), which reduce EL-induced photobleaching in *ch1* (Shumbe et al., 2016, Tano et al., 2023).

A third mutant used to understand how ^1^O_2_ signals regulate nuclear gene expression and PCD is *plastid ferrochelatase two* (*fc2*) (Woodson et al., 2015, Scharfenberg et al., 2015), which affects one of the two conserved chloroplast ferrochelatases that generate heme in plants. When grown under diurnal light cycling conditions, *fc2* mutants rapidly accumulate protoporphyrin IX (PPIX) (Woodson et al., 2015), the substrate of FC2 and a tetrapyrrole intermediate. Like Pchlide, PPIX can also produce ^1^O_2_ upon exposure to light (Duez et al., 2001). Thus, *fc2* mutants produce a burst of ^1^O_2_ at dawn when grown under cycling light conditions. This ^1^O_2_ rapidly changes nuclear gene expression (within 30 minutes) and chloroplast degradation (within three hours). After eight hours, PCD begins to occur (Woodson et al., 2015, Fisher et al., 2022). This response to ^1^O_2_ is not stage-specific, as *fc2* plants can be grown to adulthood in constant light conditions that avoid PCD. After a shift to cycling light conditions, however, *fc2* plants will exhibit PCD and growth inhibition (Woodson et al., 2015, Lemke et al., 2021, Tano et al., 2023).

Although *fc2* plants do not exhibit PCD in constant light conditions, they still accumulate low levels of ^1^O_2_ (Fisher et al., 2022). This leads to a subset of chloroplasts being degraded via the central vacuole through a process resembling fission-type microautophagy (Lemke et al., 2021, Lemke and Woodson, 2022). It is unknown how ^1^O_2_-damaged chloroplasts are recognized by the cell. Chloroplasts in *fc2* cotyledons and leaves do, however, become ubiquitinated at the chloroplast envelope, suggesting ubiquitylation as a possible mechanism for the selective turnover of ^1^O_2_-damaged chloroplasts (Woodson et al., 2015).

To gain insight into the mechanisms involved in ^1^O_2_-mediated PCD, CQC, and retrograde signaling, a forward genetic screen was performed to identify genetic suppressors of these *fc2* stress phenotypes in seedlings (Woodson et al., 2015). Here, *fc2* mutants were further mutagenized using ethyl methanesulfonate (EMS) mutagenesis and screened for secondary *fc2* suppressor (*fts*) mutations that suppress the conditional PCD phenotype of *fc2* seedlings. To date, twelve *fts* mutations have been identified and mapped to seven loci (Woodson et al., 2015, Alamdari et al., 2020, Alamdari et al., 2021). These *fts* mutations have been categorized into three classes. Class I *fts* mutants, reduce ^1^O_2_ accumulation by directly or indirectly reducing tetrapyrrole biosynthesis; Class II mutants, block ^1^O_2_ signaling without affecting chloroplast development; and Class III mutations, block ^1^O_2_ signaling with a concurrent delay in chloroplast development.

Class I mutants include those with lesions in *SUBUNIT H OF MG-CHELATASE* (*ChlH* or *GUN5*), which encodes a subunit of the Mg^2+^ chelatase that converts PPIX to Magnesium PPIX, the first intermediate of the chlorophyll branch of the tetrapyrrole pathway. Also included are mutants with lesions in *TRANSLOCON AT THE OUTER ENVELOPE MEMBRANE OF CHLOROPLASTS 159* (*TOC159*) and 33 (*TOC33*), which encode essential plastid protein import components. These mutations reduce the accumulation of ^1^O_2_ due to a reduction of tetrapyrrole synthesis (directly (*chlh*) or indirectly (*toc159*, *toc33*)) (Woodson et al., 2015). As such, they also have clear pale seedling phenotypes due to reduced levels of chlorophyll content. Although they may not represent true signaling mutants, they support the hypothesis that ^1^O_2_ accumulation leads to PCD and CQC.

The only class II mutant reported has a lesion in *PUB4*, which encodes a cytoplasmic E3 ubiquitin ligase. As *pub4-6* does not reduce chlorophyll or ^1^O_2_ levels, it is hypothesized that PUB4 likely acts downstream of the ^1^O_2_ signal, possibly by controlling the ubiquitylation of damaged chloroplasts (Woodson et al., 2015). Recent work has shown that PUB4 plays diverse roles in physiology, opening the possibility that PUB4 may target multiple proteins for ubiquitination or that chloroplast turnover pathways play a broader role in plant physiology than previously thought (Wang et al., 2017, Desaki et al., 2019, Jeran et al., 2021, Wang et al., 2022).

Class III mutants include those with lesions in *PENTATRICOPEPTIDE-REPEAT-CONTAINING PROTEIN 30* (*PPR30*), “*mitochondrial*” *TRANSCRIPTION TERMINATION FACTOR 9* (*mTERF9*) (Alamdari et al., 2020), and *CYTIDINE TRIPHOSPHATE SYNTHASE 2* (*CTPS2*) (Alamdari et al., 2021). PPR30 and mTERF9 are proteins proposed to play a role in the post-transcriptional regulation of plastid gene expression (Barkan and Small, 2014, Wobbe, 2021), while CTPS2 plays a role in maintaining dCTP levels for plastid DNA synthesis (Bellin et al., 2021). As such, all three mutations limit plastid gene expression. Like Class II mutants, Class III mutants still accumulate ^1^O_2_ in the seedling stage, suggesting they block a chloroplast ^1^O_2_ signal. Unlike Class II mutants, however, Class III mutants are pale as seedlings due to impaired chloroplast development. These mutations led to the hypothesis that a plastid-encoded product is required for the ^1^O_2_ signal.

The *fts* mutant screen only identified single recessive mutant alleles (except for *pub4-6*, which is semi-dominant) (Woodson et al., 2015). This reflects a limitation of EMS mutagenesis, which is most useful for identifying loss-of-function alleles that generally encode positive, rather than negative, regulators of signaling. To gain a more comprehensive picture of the genes and mechanisms involved in chloroplast ^1^O_2_ signaling, we aimed to identify dominant gain-of-function mutations that block these pathways. To this end, we used activation tagging, which involves randomly inserting T-DNAs into the plant genome. These sequences include *35S* enhancer elements that can overexpress nearby genes (Weigel et al., 2000).

We have used this method to identify eight dominant *fc2 activation-tagged suppressor* (*fas*) mutations that suppress chloroplast ^1^O_2_ signaling. Most of these mutations only affect ^1^O_2_ signaling in the adult stage, indicating that stage-specific pathways may exist. Furthermore, most *fc2 fas* mutants appear to specifically affect responses to ^1^O_2_, rather than general ROS responses, highlighting the specificity of our screen. However, one mutant, *fc2 fas2*, is tolerant to a wide range of abiotic stresses, suggesting that plants can also employ general stress tolerance mechanisms to overcome ^1^O_2_ stress.

## Methods

### Biological material and standard growth conditions

The wild type (wt) used in this study was *Arabidopsis thaliana* ecotype *Columbia* (Col-0). T-DNA lines GABI_766H08 (*fc2-1*) (Woodson et al., 2011) from the GABI-Kat collection (Kleinboelting et al., 2012) and SAIL_129_B07 (*atg5-1*) (Thompson et al., 2005) from the SAIL collection (Sessions et al., 2002) were described previously. The *fc2 toc159* (*fts1*), *fc2 toc33* (*fts4*), *fc2 chlH* (*fts8*), *fc2 pub4-6* (*fts29*), *fc2 ppr30-1* (*fts3*), *fc2 mterf9* (*fts32*), *fc2 ctps2-5* (*fts39*) mutants were described previously (Woodson et al., 2015, Alamdari et al., 2020, Alamdari et al., 2021). Additional information on these lines is listed in **Table S1**. Activation-tagged *fas* mutants were generated using the pSKI015 vector (Weigel et al., 2000) to transform Arabidopsis *fc2* mutants. *fc2 fas* double mutants were confirmed by extracting gDNA (following a CTAB-based protocol (Healey et al., 2014)) using PCR-based markers for the GABI_766H08 T-DNA and the pSKI015 activation tag T-DNA. Primer sequences can be found listed in **Table S2**.

Seeds, seedlings, and adult plants were germinated and grown as previously described (Woodson et al., 2015, Lemke et al., 2021) with minor deviations (most notably the use of LED plant growth chambers for adult plant growth) and detailed in the **supplemental methods** section. Unless otherwise specified, standard conditions used to grow seedlings were ∼110-115 µmol photons m^-2^ sec^-1^ at 21°C in fluorescent light chambers ((Percival^®^ model CU-36L5), set to constant light (24h light) or diurnal cycling light (6h light/18h dark) conditions) and standard conditions used to grow adult plants were ∼110-120 µmol photons m^-2^ sec^-1^ at 21°C with 60% humidity in an LED plant growth chamber ((Hettich PRC 1700), set to constant light (24h light) or diurnal cycling light (16h light/8h dark) conditions). *Escherichia coli* and *Agrobacterium tumefaciens* were grown in liquid Miller nutrient broth or solid medium containing 1.5% agar (w/v). Cells were grown at 37°C (*E. coli*) or 28°C (*A. tumefaciens*) with the appropriate antibiotics.

### Mutagenesis by activation-tagging

The pSKI015 vector (containing *35S* enhancers and a Basta resistance marker gene) was transformed into the *A. tumefaciens* strain GV3101, which was subsequently used to transform *fc2* mutants via the Agrobacterium-mediated floral dip method (Clough and Bent, 1998). T_1_ plants were selected on Basta-soaked soil (1.5ml of Bayer Finale herbicide per 2 liters of H_2_O per flat) and screened for suppression of the *fc2* PCD phenotype in diurnal cycling light (16h light/8h dark) conditions in a fluorescent light growth chamber (Percival model AR-75LX), ∼80-100 µmol photons m^-2^ sec^-1^ at 21°C). T_1_ suppressor candidates were propagated (an estimated 11,027 T_1_ plants were screened, and 736 were selected for further monitoring). T_2_ lines were monitored for a robust suppressor phenotype (suppression of PCD) in cycling light and co-segregation of this phenotype with Basta resistance. Finally, homozygous *fas* lines were selected for further testing in the T_3_ generation based on 100% Basta-resistance and a robust *fc2* suppressor (*fas*) phenotype. We isolated a total of seven *fas* mutants (*fc2 fas1*, *fc2 fas2*, *fc2 fas3*, *fc2 fas4*, *fc2 fas6*, *fc2 fas7*, *fc2 fas8*, *fc2 fas9*) that passed these stringent criteria and two that did not (*fc2 fas3* and *fc2 fas5*). In the case of *fc2 fas3* and *fc2 fas5*, the suppressor phenotype did not co-segregate with Basta resistance. As such, PCR primers (**Table S2**) were designed to probe for the presence of the pSKI015 TDNA. An intact pSKI015 TDNA was observed to be present in all *fas* mutants except for *fc2 fas3* and *fc2 fas5*. The *fas* phenotype was also observed to co-segregate with altered rosette morphology in *fc2 fas2*, *fc2 fas4*, and *fc2 fas7*, suggesting a linkage between these phenotypes in these mutants.

Finally, these *fas* mutants were tested for dominance of the *fas* phenotype. Here, *fc2* was fertilized as the maternal line using pollen from each respective *fc2 fas* mutant as the paternal line (♀*fc2* x ♂*fas*). The F_1_ generation plants were then grown in cycling light stress conditions alongside their parent plants to monitor for *fas* phenotypes. The *fas* phenotypes of *fc2 fas1*, *fc2 fas2*, *fc2 fas3*, *fc2 fas4*, *fc2 fas6*, *fc2 fas7*, *fc2 fas8*, *fc2 fas9* were recapitulated in the F_1_ generation, confirming their dominance. Although an intact pSKI015 TDNA was no longer detected in *fc2 fas3*, we included it in this study as it has a dominant phenotype. Consequently, *fc2 fas3* likely still contains a mutation generated during the screen, possibly an incomplete T-DNA or an insertion/deletion caused by the excision of the T-DNA. The *fc2 fas5* mutant did not have a dominant phenotype and lacked the pSKI015 TDNA. Thus, we excluded it from further testing.

### Plant growth hormone and abiotic stress treatments

#### Exogenous GA_3_ treatment

Plants were grown in constant light conditions for seven days and then transplanted to soil. Beginning at ten days, plants were sprayed with 1 mL of 10^-4^ M GA_3_ (suspended in H_2_O) every two days, as described in (Ribeiro et al., 2012). For a mock treatment, untreated plants were sprayed with 1 mL of H_2_0. At 14 days, plants were transferred to cycling light conditions (16h light, 8h dark) for seven days or left in constant light conditions. Physiological observations and measurements were taken at 21 days.

#### Abiotic stress tests

For all abiotic stress tests, plants were grown in constant light LED chambers for 21 days (EL, MV, heat, and freezing treatments) or 23 days (dark-induced carbon starvation) and then subjected to various stresses. Chlorophyll fluorescence measurements were taken at least three times at the beginning of stress treatment. The starting F_v_/F_m_ values of each genotype consistently started within the F_v_/F_m_ range expected for unstressed plants (∼0.81-0.84), except *fc2 toc33*, which consistently had a significantly reduced starting F_v_/F_m_ range of ∼0.78-0.81 **(Figs. S1a-c)**.

#### Excess light stress

Excess light (EL) treatments used 21-day-old plants, which were exposed to 1450-1550 µmol photons m^-2^ sec^-1^ white light at 10 °C (to offset for excess heat generated by the EL panel) for 24 hours in a Percival LED 41L1 chamber (with SB4X All-White SciBrite LED tiles). The average leaf temperature from 6 representative plants (after 2 hours in EL conditions) was 19.7 °C, which was measured using an Etekcity Lasergrip 630 Infrared Thermometer. Chlorophyll fluorescence measurements of the same plants were collected before treatment and after 6h and 24h of EL treatment. Lesion formation was assessed in the same plants immediately after 24h EL treatment (22-day-old plants), prior to any regreening and/or new growth during recovery. After 24h of EL, plants were returned to a constant light LED chamber and allowed to recover for three days, at which point representative plants were imaged (25-day-old plants).

#### Methyl Viologen stress

MV treatment (20 µM or 200 µM) was applied to 21-day-old plants grown in constant light conditions. MV (Sigma-Aldrich) was dissolved in H_2_O to create 20 µM or 200 µM solutions, which were applied to plants via a generic perfume atomizer (4 sprays each, approximately 0.5 mL per plant). Upon treatment, plants were returned to their respective growth chambers for 24 hours, at which point chlorophyll fluorescence was measured (22-day-old plants). Plants were subsequently returned to a constant light LED chamber and allowed to recover for three days. After recovery (and time allowed for MV-induced lesion formation), lesions were counted, and representative plants were imaged (25-day-old plants).

#### Heat stress

Heat stress was applied by placing 21-day-old plants in a 40 °C incubator (in the dark) for 16 or 24 hours. For no heat control, a dome was placed over a flat of 21-day-old plants, wrapped in foil, and placed in the dark for 24 hours at 21°C. Plants were allowed to cool on a laboratory bench in dim light (10-15 µmol photons m^-2^ sec^-1^) for one hour, and then chlorophyll fluorescence was subsequently measured (22-day-old plants). Plants were then returned to constant light conditions and allowed to recover for three days, at which point representative plants were imaged (25-day-old plants).

#### Freezing stress

Freezing stress was applied by incubation at −20°C in a freezer. To account for the possible unequal insulating effect of dry soil or uneven amounts of water saturation, soil pots of 21-day-old plants were weighed individually, and their weights were equalized by adding water to the soil. Pots were then returned to their flats and incubated at 4°C (cold acclimated set) or at 21°C (unacclimated) in the dark for 16 hours. One flat from each condition was then transferred to −20°C for 1 hour. Flats were then removed from the freezer and allowed to thaw for two hours on a laboratory bench in dim light, at which time chlorophyll fluorescence of plants was measured (22-day-old plants). Afterward, the plants were returned to constant light conditions for three days, at which point representative plants were imaged (25-day-old plants).

#### Carbon starvation

Carbon starvation was achieved by moving 23- or 21-day-old plants into the dark at 21°C for five or seven days, respectively. At the end of each treatment regimen, plants were left on a laboratory bench in dim light to equilibrate for one hour before chlorophyll fluorescence measurements were taken (28-day-old plants). Plants were then returned to standard constant light conditions to recover for four days, at which point representative plants were imaged (32-day-old plants). All abiotic stress tests were conducted at least three times with consistent results, and representative experiments are shown.

#### Confirmation of T-DNA mutant lines by PCR genotyping

Genotyping Primers were designed using the SIGnAL (http://signal.salk.edu/) T-DNA primer design tool or Primer3 https://www.primer3plus.com/ (Untergasser et al., 2012) **(Table S2)**. Primers for *fc2-1* were used to confirm the background genotype of *fas fc2* candidates. Primers were also designed to probe for different regions of the pSKI015 TDNA in *fas fc2* candidates. Here, pSKI015-790F/pSKI015-1974R, pSKI015-2261F/pSKI015-2972R, and pSKI015-3772F/pSKI015-4962R were used to probe different regions of the pSKI015 T-DNA. All PCR was performed as described (Lemke et al., 2021) and detailed in the **supplemental methods** section.

#### Biomass measurements

Biomass was assessed as previously described (Lemke et al., 2021) and detailed in the **supplemental methods** section.

### Chlorophyll fluorescence measurements

Chlorophyll fluorescence measurements were conducted as previously described (Lemke et al., 2021) and detailed in the **supplemental methods** section.

### Chlorophyll measurements

Plant chlorophyll content was measured as previously described (Woodson et al., 2015) and detailed in the **supplemental methods** section.

### Cell death measurements

Cell death was assessed using trypan blue staining as previously described for seedlings (Alamdari et al., 2020) and detailed in the **supplemental methods** section.

### RNA extraction and Reverse transcription-quantitative polymerase chain reaction (RT-qPCR)

Total RNA was extracted from plants, cDNA was synthesized, and RT-qPCR was conducted as previously described (Lemke et al., 2021) and detailed in the **supplemental methods** section. All expression data were normalized according to *ACTIN2* expression. All primers used for RT-qPCR are listed in **Table S2**.

#### RT-qPCR data visualization heatmaps

A data matrix reflecting the RT-qPCR %wt change was compiled for each list. The data used for these heatmaps, and their corresponding significance can be found in **(Table S3-6)**. Heatmaps were generated using the heat mapper package (https://github.com/WishartLab/heatmapper) on http://www.heatmapper.ca/ (Babicki et al., 2016).

### Singlet oxygen measurements

^1^O_2_ was measured with Singlet Oxygen Sensor Green dye (SOSG, Molecular Probes 2004) using a protocol adapted from (Alamdari et al., 2020) and optimized for adult leaf tissue. Here, plants were grown in standard constant light conditions for 21 days and then transferred to diurnal cycling light conditions (16h light, 8h dark) for two days. At the end of the second day, at least 12 leaf disks (4 mm) were cut (using a cork borer) from true leaves (#’s 3-6) selected from separate plants and placed in 50 mM phosphate buffer, pH 7.5 (250 ul in 1.5 ml tubes), which were then wrapped in foil and returned to the growth chamber overnight (16 hours). One hour before light exposure (subjective dawn) on day three, 100 µg of SOSG was dissolved in 30 μl of 100% MeOH. 70μl of 50 mM phosphate buffer, pH 7.5 + 0.1% tween 20, was added to the SOSG solution (final concentration 1 μg/ul SOSG solution). 45 μl of SOSG solution was added to the leaf disk tubes in a dark room illuminated with dim green light and then placed in a domed desiccator wrapped two times in aluminum foil. Leaf disks were then vacuum infiltrated (∼ −25 mmHg) in the dark for 30 minutes, then incubated for another 30 minutes. Leaf disks were then removed from the dark and returned to the growth chamber in light for 30 min. Leaf disks were washed twice with 1 mL 50 mM phosphate buffer, pH 7.5, and returned to the chamber for 1.5 hours. Leaf disks were then imaged, and SOSG fluorescence was measured with a Zeiss Axiozoom 16 fluorescent stereo microscope equipped with a Hamamatsu Flash 4.0 camera and a GFP fluorescence filter. The average SOSG signal (fluorescence per mm^2^) of each leaf disk was quantified using ImageJ. Experiments were conducted in two separate groups: Group A (wt, *fc2*, *fts4*, *fts29*, *fc2 fas1*, *fc2 fas2*, and *fc2 fas3*) and Group B (wt, *fc2*, *fc2 fas4*, *fc2 fas6*, *fc2 fas7*, *fc2 fas8*, and *fc2 fas9*). Leaf disks from wt and *fc2* were also treated with SOSG but left in the dark (-light) to assess the specificity of the signal we were measuring. Any SOSG (-light) signal was negligible relative to SOSG (+light) leaf disks, being only 5.2% of SOSG (+light) in wt and 1.8% of SOSG (+light) in *fc2*. Leaf disks from wt and *fc2*, without SOSG treatment, were analyzed to account for possible background fluorescence of plant leaf tissue. Any autofluorescence observed was negligible relative to SOSG-treated leaf disks, 0.0005 % of SOSG (+light) in wt and 0.000001% of SOSG (+light) in *fc2*. The results of this assay development and optimization can be found in **Fig. S2**.

### Graphical model creation

**Fig. S3** was created using online BioRender software (https://biorender.com/).

## Results

### Defining the selection criteria for the identification of mutations that suppress fc2 phenotypes in the adult stage

To identify new dominant gain-of-function mutations that suppress ^1^O_2_-induced stress responses, we turned to activation tagging, which allows us to screen plants for suppressor phenotypes in the T1 generation. The advantage of this strategy is that all mutants are in a hemizygous state. If they suppress conditional *fc2* phenotypes (e.g., do not exhibit PCD in cycling light conditions), they must be dominant alleles. However, this necessitates that the plants be screened as adults after being selected for transgenesis. It has been previously shown that when grown as adults in cycling light conditions, *fc2* mutants do initiate ^1^O_2_ signaling (Woodson et al., 2015, Lemke et al., 2021, Tano et al., 2023). However, *fts* suppressor mutants have mostly been characterized as seedlings and it is unclear how they behave as adults (a summary of published *fts* mutants and their seedling phenotypes can be found in **Table S7)**. With this in mind, it was important first to determine if performing a genetic screen with adult *fc2* plants can allow for identifying new suppressor mutations.

To this end, wt and *fc2* plants along with Class I (*fc2 toc159*, *fc2 toc33*, and *fc2 chlH)*, Class II (*fc2 pub4-6*), and Class III (*fc2 ppr30, fc2 mterf9,* and *fc2 ctps2*) suppressor mutants were grown for 21 days in constant light (24h) conditions or for 14 days in constant light conditions and then transferred to diurnal cycling light (16h light/8h dark) conditions for seven days. As previously reported (Woodson et al., 2015), *fc2* appears healthy (no leaf lesions) under constant light conditions **(Fig. S4a)**. Under these conditions, all *fc2 fts* mutants also appear healthy, but *fc2 toc159*, *fc2 toc33,* and *fc2 mterf9* exhibit pale and chlorotic phenotypes relative to *fc2*. After 7 days in cycling light conditions, *fc2* mutants form necrotic lesions while wt does not **(Fig. S4a)**. Class I and Class II suppressor mutants appear to suppress the formation of these lesions, while Class III mutants still accumulate some lesions. When trypan blue stains were conducted to measure cell death in plants grown under cycling light conditions, *fc2* exhibited significantly more cell death than wt (**Figs. 4b and c**). Compared to *fc2*, the Class I and Class II suppressor mutants all had significantly reduced trypan blue staining in leaves. On the other hand, Class III suppressor mutants still exhibited trypan blue staining and were not significantly different from *fc2*.

Next, total chlorophyll content was measured in plants grown in constant light conditions. Here, *fc2* appeared to have lower levels of total chlorophyll than wt, but not to a significant extent **(Fig. S4d)**. Under the same conditions, *fc2 toc159*, *fc2 toc33* and *fc2 mterf9* had significantly less chlorophyll than *fc2*, while *fc2 pub4-6, fc2 chlH*, *fc2 ppr30*, and *fc2 ctps2* did not. Thus, we established suppressor mutant classification in the adult stage: Class I suppressor mutants significantly block PCD and generally have reduced levels of chlorophyll, Class II suppressor mutants significantly block PCD and do not have reduced levels of chlorophyll, and Class III suppressor mutants do not significantly block PCD and generally do not have reduced levels of chlorophyll (**Table S8**). These results show that Class I and II mutant phenotypes generally translate to the adult stage. As Class II suppressor mutants are expected to affect signaling components, we chose to prioritize their identification, which is easily distinguished by a lack of lesions and chlorosis.

### An activation tagging screen to identify dominant suppressors of the fc2 programmed cell death phenotype

The above results suggested that the adult *fc2* plants can be screened for suppression of ^1^O_2_-mediated PCD signals. As such, we transformed *fc2* plants with the pSKI015 activation tagging vector, which contains a transfer DNA (T-DNA) containing four repeats of the cauliflower mosaic virus (CaMV) *35S* enhancer sequence (Weigel et al., 2000) **(Fig. S3a)**. This can lead to the overexpression of proximal genes **(Fig. S3b)**. Consequently, this enhanced expression is subject to native tissue- and stage-specific transcriptional programming, thereby increasing the probability of identifying biologically relevant genes, as opposed to constitutive ectopic expression by the full CaMV *35S* promoter (Benfey et al., 1990). Activation-tagged *fc2* transformants (hereafter referred to as mutants) were selected in the T_1_ generation and grown under diurnal light cycling conditions (16h light/8h dark) **(Fig. S3c)**. We then chose candidates in the T_1_ generation that appeared to reduce the number or size of lesions in true leaves caused by ^1^O_2_. We screened an estimated 11,000 T_1_ transformants and chose 736 for more stringent screening in the T_2_ generation. This led to the identification of nine potential *fas* mutants that block the *fc2* PCD phenotype. To confirm the dominance of these mutations, all *fc2 fas* mutants were backcrossed to the *fc2* parent, and the phenotypes of the F_1_ generation were tested. The phenotypes of *fc2 fas1*, *fc2 fas2*, *fc2 fas3*, *fc2 fas4*, *fc2 fas6*, *fc2 fas7*, *fc2 fas8*, *fc2 fas9* were recapitulated in the F_1_ generation, confirming their dominance **(Fig. S5a)**. Trypan blue stains were conducted to measure cell death. All included *fc2 fas* mutants showed a significant reduction of PCD, relative to *fc2*, in the F_1_ generation **(Figs. S5b and c)**. The *fc2 fas5* mutant (not pictured) did not have a dominant phenotype and was excluded from further testing.

### Activation-tagged fas fc2 mutants suppress cell death under light cycling conditions

We first investigated the visual phenotype of our eight confirmed *fc2 fas* mutants under permissive (constant light conditions) and stress (14 days of constant light conditions and then moved to cycling light conditions (16h light/8h dark) for 7 days) (**Fig. 1a**). Under constant light conditions, *fc2* mutants appear healthy without an obviously altered rosette morphology, but are visually smaller than wt. When exposed to cycling light conditions, however, *fc2* mutants form necrotic lesions. We also included *fts* mutants *toc33 fc2* (Class I suppressor mutant) and *pub4-6 fc2* (Class II suppressor mutant) as controls for different forms of *fc2* PCD suppression. Under the same cycling light conditions, all *fts* and *fas* mutations suppress the conditional PCD observed in *fc2* (**Fig. 1a**). When trypan blue stains were conducted to assess cell death, all *fc2 fts* and *fc2 fas* mutants had significantly less PCD compared to *fc2* (**Figs. 1b and c**). These data confirm that all *fas* mutations suppress the conditional *fc2* PCD phenotype in the adult stage.

**Figure 1.**
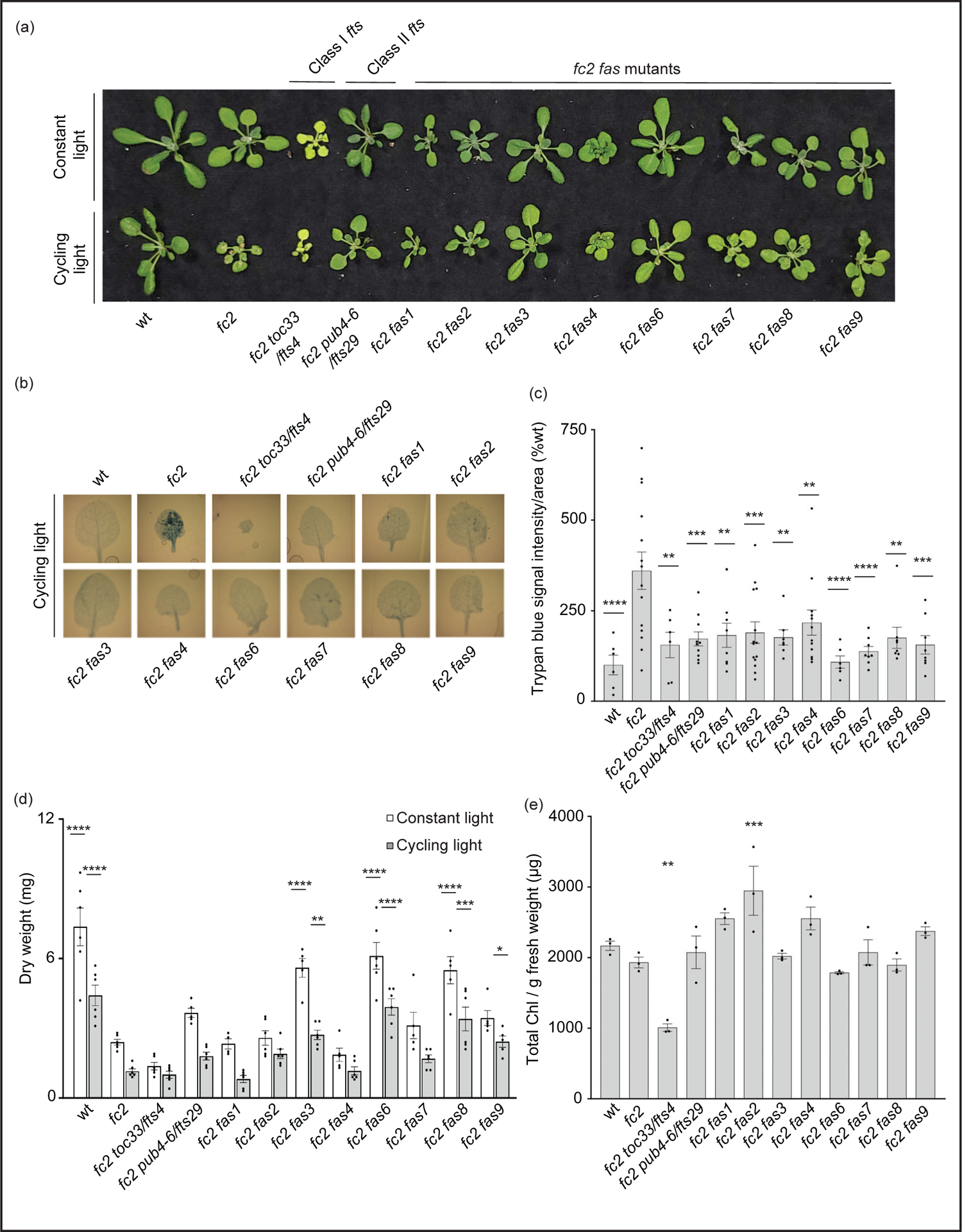
*fas* mutations suppress light-induced programmed cell death in *fc2* Eight dominant *fc2 fas* mutants were assessed for their capacity to suppress the conditional programmed cell death phenotype of *fc2*. Plants were grown for 21 days in constant light conditions or 14 days in constant light conditions and then transferred to cycling light conditions (16h light/8h dark) for 7 days. (a) A photograph of representative plants from both light conditions. (b) Representative images of leaves collected from plants grown under cycling light conditions and stained with trypan blue. The dark blue color indicates dead tissue. (c) Quantification of trypan blue stains shown in b (n ≥ 6 leaves from individual plants). (d) Dry weight biomass (mg) of plants (aerial tissue) grown under constant light and cycling light conditions (n ≥ 4 plants). (e) The total chlorophyll content (μg/g fresh weight (FW)) of plants grown in constant light conditions (n = 3 leaves from individual plants). Trypan blue stain, biomass, and total chlorophyll content quantification were tested using a One-way ANOVA, and Dunnett’s multiple comparisons post hoc was used to test variation between genotypes relative to wt (* = *P* ≤ 0.05, ** = *P* ≤ 0.01, *** = *P* ≤ 0.001, **** = *P* ≤ 0.0001). Error bars = +/- SEM. Closed circles indicate individual data points.

We next measured the dry-weight biomass of aerial tissue from the same plants. When grown under constant light conditions, *fc2* was observed to have reduced biomass relative to wt (**Fig. 1d**). This reduction in biomass relative to wt is even more pronounced in *fc2* exposed to cycling light conditions. Under both constant light and cycling light conditions, *fc2 fas3*, *fc2 fas6*, and *fc2 fas8* were observed to have a biomass greater than *fc2* in each respective condition, whereas *fc2 fas9* had a greater biomass than *fc2* only in cycling light conditions. Conversely, *fc2 fas1*, *fc2 fas2*, *fc2 fas4,* and *fc2 fas7* were not observed to have significantly greater biomass than *fc2* in either constant light or cycling light conditions.

We also observed no *fc2 fas* mutants to be obviously pale when grown in constant light conditions (**Fig. 1a**). To assess chlorophyll levels, we measured the total chlorophyll content of these plants. As expected, *fc2 toc33* had significantly reduced chlorophyll compared to *fc2*, while *fc2 pub4-6* did not. Finally, none of the *fc2 fas* mutants had significantly reduced chlorophyll levels compared to *fc2* (**Fig. 1e**). We did, however, observe increased chlorophyll content in *fc2 fas2* compared to *fc2*. These data suggest that *fas* mutations do not lead to reduced chlorophyll levels in the adult stage, and, as such, are unlikely to be Class I suppressor mutations.

As our dominant *fas* mutants have a PCD suppressor phenotype in the adult stage (21 days old), we next asked if suppression of the *fc2* phenotypes is also active at the seedling stage. Under cycling light conditions (6h day/18h night), wt seedlings green normally, whereas *fc2* seedlings bleach and die (**Fig. 2a**). As expected, the *fc2 fts* mutants (*fc2 toc33* and *fc2 pub4-6 fts*) suppress this PCD phenotype. Among the *fc2 fas* mutants, *fc2 fas3* and *fc2 fas7* were the only ones observed to visibly suppress PCD at the seedling stage. This pattern of cell death was confirmed via trypan blue stains (**Figs. 2b and c**). When grown under constant light conditions, no *fas* mutants exhibited a visibly pale phenotype (**Fig. 2a**), or significantly reduced chlorophyll content (**Fig. 2d**) compared to *fc2*. Together, these data demonstrate that the *fas3* and *fas7* mutations are the only *fas* mutations that suppress the conditional *fc2* PCD phenotype in the seedling stage, illustrating that the other six *fas* mutations have a stage-specific effect on stress signaling. Furthermore, the limited changes of chlorophyll content in the eight *fc2 fas* mutants in the adult and seedling stages suggest that they are not Class I or III suppressor mutants.

**Figure 2.**
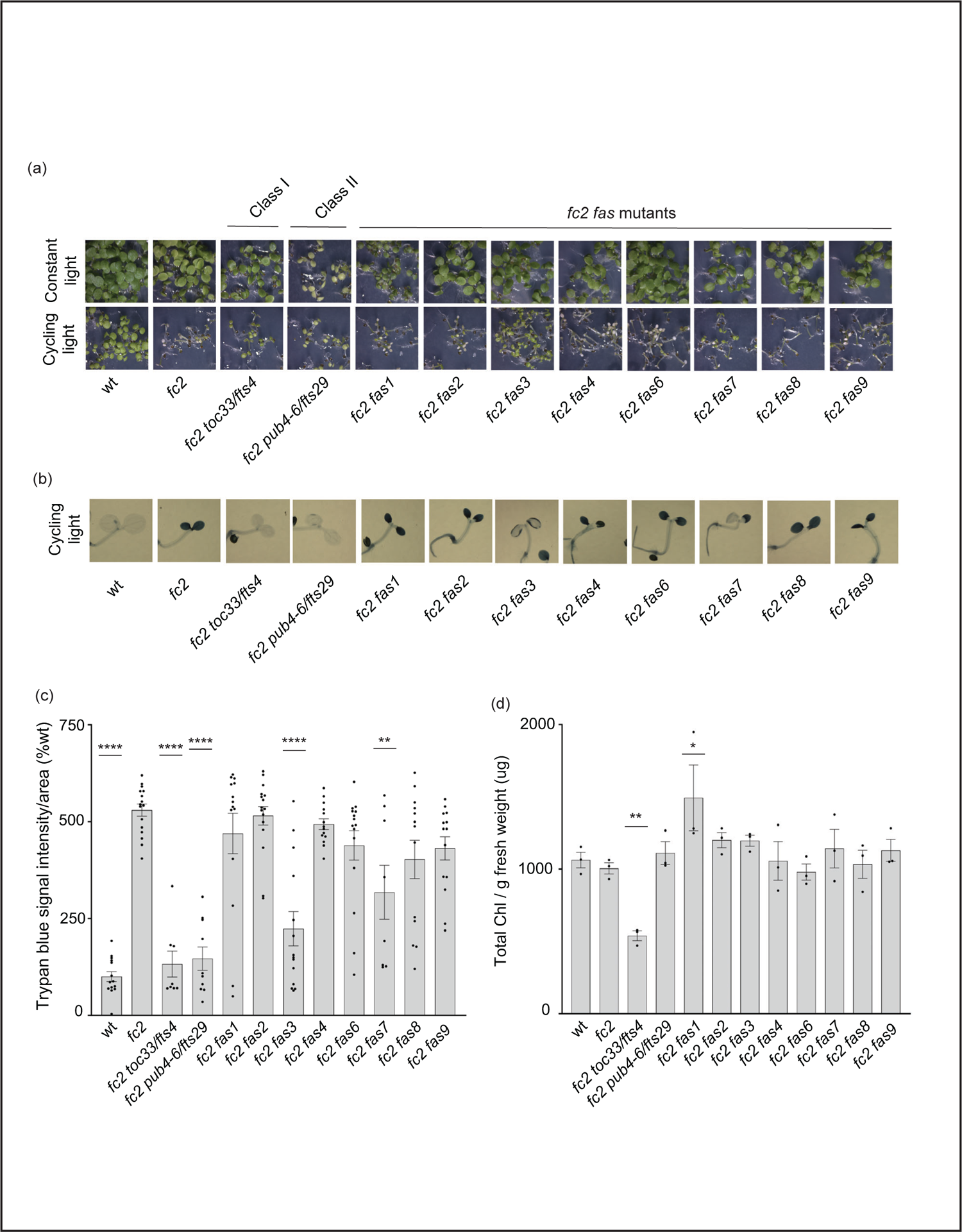
Assessment of seedling stage phenotypes of *fc2 fas* mutants Suppression of cell death by the *fc2 activation tagged suppressor* (*fas*) mutations was assessed in the seedling stage. Seedlings were grown for seven days in constant light (24h) or cycling light (6h light/ 18h dark) conditions. (a) Image showing seeding phenotypes of plants grown under constant or cycling light conditions. (b) Trypan blue stains of seedlings grown under cycling light conditions. The darker blue stain indicates dead tissue (c). Quantification of trypan blue stains shown in b (n ≥ 6 biological replicates). (b) Quantification of chlorophyll content (μg/g fresh weight) of seedlings grown in constant light (n = 3 biological replicates). Trypan blue stain and chlorophyll content quantification were tested using a one-way ANOVA, and a Dunnett’s multiple comparisons post hoc was used to test variation between genotypes relative to *fc2* (* = *P* ≤ 0.05, ** = *P* ≤ 0.01, **** = *P* ≤ 0.0001). Error bars = +/- SEM. Closed circles indicate individual data points.

### fas mutations affect singlet oxygen-induced retrograde signaling

In addition to PCD, chloroplast-generated ^1^O_2_ activates retrograde signals to regulate nuclear gene expression (op den Camp et al., 2003, Ramel et al., 2013, Woodson et al., 2015). Therefore, we next tested if retrograde signaling is affected by *fas* mutations. To this end, we probed response marker genes via RT-qPCR in 21-day-old plants grown in cycling light conditions (14 days of constant light conditions and moved to 7 days of cycling light conditions (16h light/8h dark)). These include marker genes that are induced in *fc2* seedlings under cycling light stress (Woodson et al., 2015); *HSP26.5*, *HSP22*, and *SIB1*, SORGs (op den Camp et al., 2003); *BAP1*, *ATPase*, and *NOD*, general oxidative stress response genes (Baruah et al., 2009); *ZAT12*, *CYC81D8* and *GST*, and H_2_O_2_ response genes (Laloi et al., 2007); *APX1* and *FER1*.

We started with marker genes identified to be induced in *fc2* seedlings grown under cycling light conditions (*fc2*-stress). Here, only *HSP26.5* and *SIB1* were significantly induced in *fc2* (relative to wt), suggesting that some variation exists in the *fc2* stress response to cycling light between the seedling and adult stages (**Figs. 3a and b**). *SIB1* expression was observed to be reduced to wt levels in *fc2 fas* mutants, except *fc2 fas4*. It should be noted that *SIB1*, a chloroplast-associated transcriptional regulator of nuclear gene expression, was also later identified as a *SORG* (Dogra et al., 2017). *HSP26.5 and HSP22* were also observed to be induced in *fc2 fas1* and *fc2 fas7*, compared to wt, respectively.

**Figure 3.**
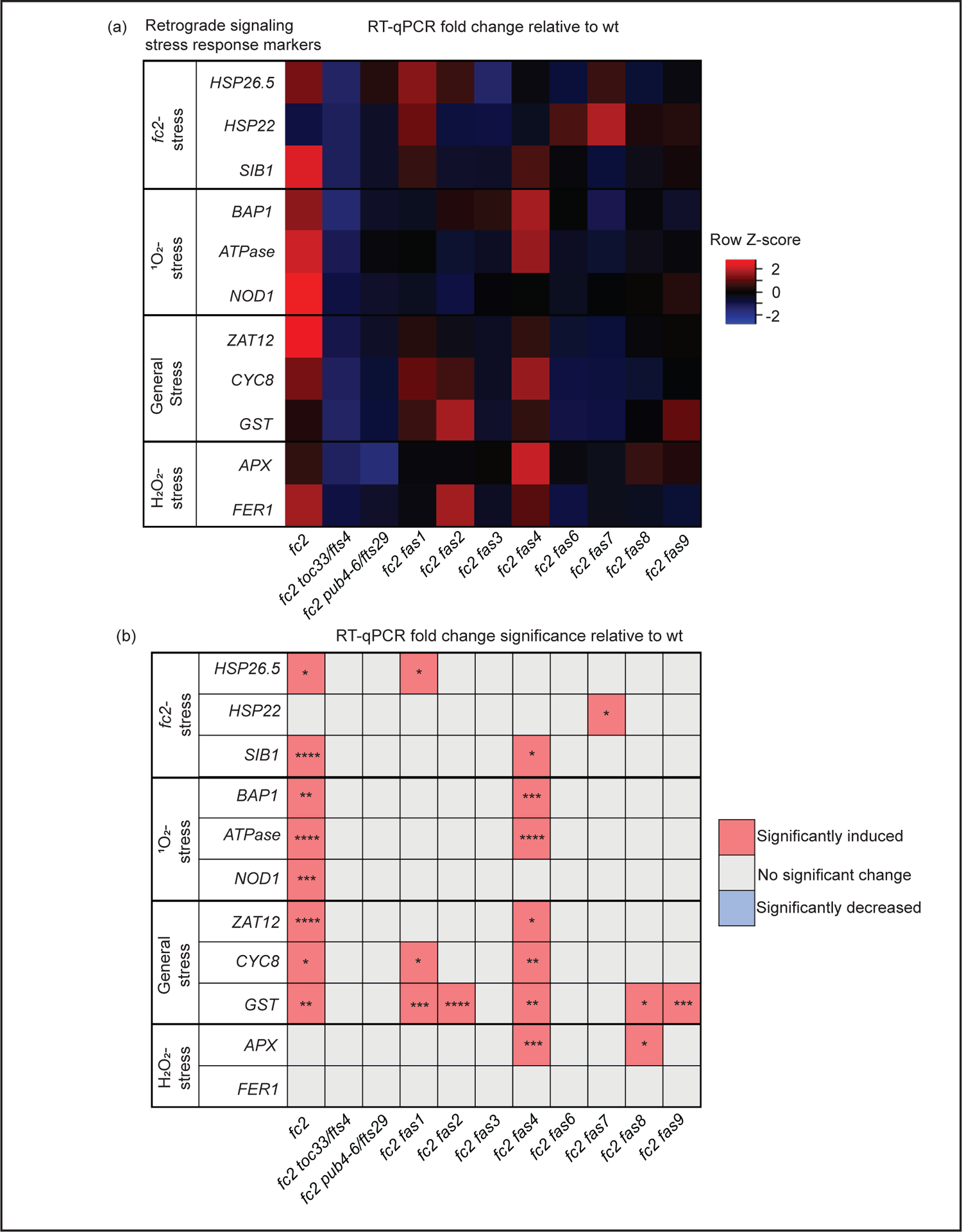
Analysis of reactive oxygen species retrograde signaling in *fc2 fas* mutants Reactive oxygen species (ROS) stress marker genes were probed via RT-qPCR to investigate the potential influence of *fas* mutations on chloroplast retrograde signaling. Transcript abundance was monitored in 21-day-old plants grown for 14 days in constant light conditions and then transferred to cycling light (16h light/8h dark) conditions for 7 days. Marker genes for *fc2*, singlet oxygen (^1^O_2_), general oxidative, and hydrogen peroxide (H_2_O_2_) stress were selected to probe for transcriptional responses via RT-qPCR. (a) A heatmap table summarizing the fold-change of transcript abundance relative to wt. Shades of red indicate increased transcript abundance, and shades of blue indicate decreased transcript abundance. (b) Table reporting significance of difference in marker transcript abundance, relative to wt. All quantification of qPCR analyses were tested using a one-way ANOVA. A Dunnett’s multiple comparisons post hoc were used to compare variation between genotypes relative to wt (* = *P* ≤ 0.05, ** = *P* ≤ 0.01, *** = *P* ≤ 0.001, **** = *P* ≤ 0.0001). n = 3 biological replicates.

As *fc2* mutants are ^1^O_2_ overproducers, we next probed *SORG* marker genes. These three genes showed significant induction in stressed *fc2* (relative to wt) (**Figs. 3a and b**). The three SORGs were no longer observed to be induced (relative to wt) in all *fc2 fas* mutants, except for *fc2 fas4*, which still retained a significant induction of *BAP1* and *ATPase*, compared to wt.

We next probed general oxidative stress response genes, which were all significantly induced in *fc2* relative to wt (**Figs. 3a and b**). *ZAT12* expression was reduced to wt levels in all *fc2 fas* mutants, except *fc2 fas4*. *CYC8* expression was decreased to wt levels in all *fc2 fas* mutants except *fc2 fas1* and *fc2 fas4*. Finally, *GST* expression was only reduced to wt levels in *fc2 fas3*, *fc2 fas6* and *fc2 fas7*.

To test if these retrograde signals and PCD were explicitly caused by ^1^O_2_, we also probed H_2_O_2_ marker genes. Neither *APX1* nor *FER1* was significantly induced in *fc2* at the adult stage (relative to wt (**Figs. 3a and b**)). However, *APX1* expression was increased in *fc2 fas4* and *fc2 fas8*, compared to wt. *FER1*, on the other hand, was not increased in any of the *fc2 fas* mutants. These data suggest that ^1^O_2_ is the predominant ROS-stress signal involved in adult cycling light-stressed *fc2,* consistent with observations made in *fc2* seedlings (Woodson et al., 2015, Alamdari et al., 2020). Except for *fc2 fas4*, all *fc2 fas* mutants reverse the induction of stress marker genes.

### *Measuring singlet oxygen accumulation in* fc2 fas *mutants*

As all *fas* mutations reduce ^1^O_2_ signaling responses in *fc2*, we next tested if ^1^O_2_ is still accumulating in these plants. To this end, we measured ^1^O_2_ accumulation in leaves from 24-day-old plants subjected to 3 days of cycling light conditions (16h light/ 8h dark) using Singlet Oxygen Sensor Green (SOSG), a dye that fluoresces in the presence of ^1^O_2_. When infiltrated with SOSG and subjected to light, *fc2* leaf disks accumulated significantly higher fluorescence levels than wt leaf disks **(Fig. S2)**. This response was light-dependent. When wt and *fc2* leaf disks were treated with SOSG but not subjected to light, lower fluorescence levels were observed, with no significant difference between genotypes. Without SOSG, virtually no fluorescence was detected in any leaf discs. Together, these results demonstrate that SOSG can be used to measure bulk ^1^O_2_ in leaf discs and that the *fc2* mutant still accumulates increased levels of ^1^O_2_ in true leaves.

We repeated this assay with the *fc2 fts* and *fc2 fas* mutants. Again, *fc2* accumulated significantly more bulk ^1^O_2_ than wt (**Figs. 4a and b**). ^1^O_2_ was also observed to accumulate in *fc2 toc33* and *fc2 pub4-6* compared to wt. The high level of ^1^O_2_ in *fc2 toc33* was unexpected as this mutant has low ^1^O_2_ levels as a seedling (Woodson et al., 2015). When we measured ^1^O_2_ accumulation in the *fc2 fas* mutants, we observed ^1^O_2_ accumulation at significantly increased levels (relative to wt) in *fc2 fas1*, *fc2 fas3, fc2 fas6*, *fc2 fas7*, *fc2 fas8*, and *fc2 fas9*. ^1^O_2_ levels in *fc2 fas4* were observed to be increased (and statistically similar to *fc2*), but were not significantly different from wt. No significant accumulation of ^1^O_2_ was observed in the *fc2 fas2* mutant compared to wt. These data suggest that most *fas* mutations do not reduce bulk ^1^O_2_ levels and, therefore, may reduce PCD and retrograde signaling by blocking a ^1^O_2_-mediated signal.

**Figure 4.**
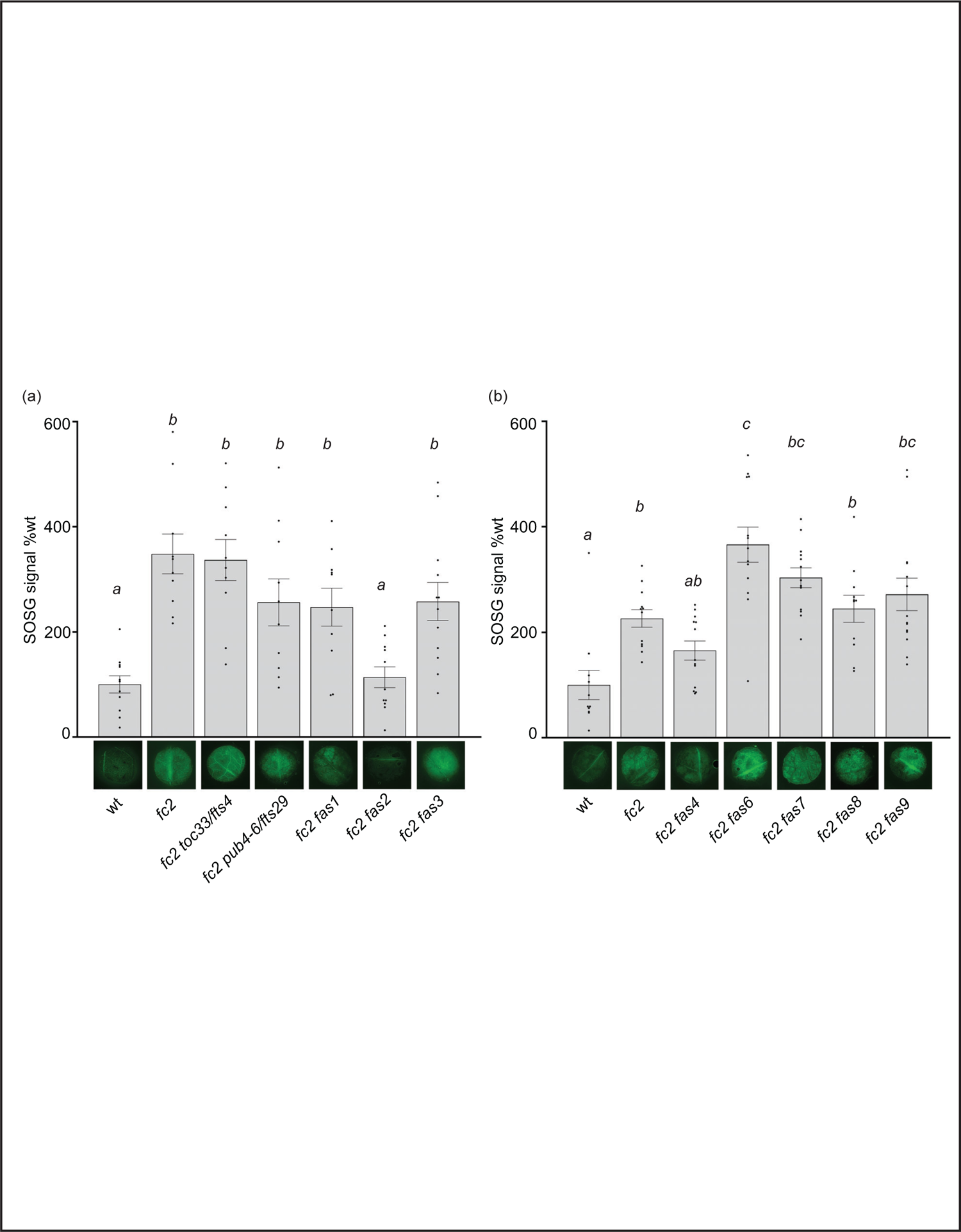
Singlet oxygen accumulation in *fc2 fas* mutants Singlet oxygen (^1^O_2_) accumulation was monitored in 24-day-old leaf tissue. Plants were grown for 21 days in constant light conditions and then transferred to cycling light conditions (16h light/8h dark) for 3 days. Leaf disks were collected and infiltrated with singlet oxygen green (SOSG). (a-b) Images of representative leaf disks showing SOSG fluorescence positioned under graphs quantifying the SOSG signal. Each panel represents an independent experiment. Brighter green fluorescence indicates higher bulk levels of ^1^O_2_. Quantification of the SOSG signal was tested with a one-way ANOVA with Tukey’s multiple comparisons post hoc to compare variation between genotypes. Different letters above bars indicate significant differences within data sets (*P* ≤ 0.05). n ≥ 10 leaf disks from replicate plants. Error bars = +/- SEM. Closed circles indicate individual data points.

### Growth hormone responses are perturbed in fc2 fas mutants

While four *fc2 fas* mutants (*fc2 fas3*, *fc2 fas6*, *fc2 fas7*, *fc2 fas8*, and *fc2 fas9*) do reverse the impaired growth rate of cycling light stressed *fc2*, another four of the *fc2 fas* mutants (*fc2 fas1, fc2 fas2*, *fc2 fas4*, and *fc2 fas7)* do not (**Figs. 1a and d**). This variability in growth rate between the different *fas* mutants led us to question whether growth hormone signaling is affected in these lines. To this end, we designed qPCR probes for key Gibberellic Acid (GA), Brassinosteroid (BR), Auxin (IAA), and Cytokinin (CK) response marker genes. We probed these growth hormone marker genes via RT-qPCR in 21-day-old plants grown in cycling light conditions (14 days of constant light conditions and moved to cycling light conditions (16h light/8h dark) for 7 days).

For GA response markers, we targeted two genes that have been reported to respond to GA: *GAST1 PROTEIN HOMOLOG 1* (*GASA1*) and *GIBBERELLIN 3-OXIDASE 1* (*GA3OX1*) (Herzog et al., 1995, Igielski and Kepczynska, 2017). We observed no differential expression pattern between wt and *fc2* (**Figs. 5a and b**). Two *fas* mutants (*fc2 fas2* and *fc2 fas7*) showed significant induction of *GASA1* (relative to wt), whereas only *fc2 fas2* showed induction of *GA3OX1* relative to wt.

**Figure 5.**
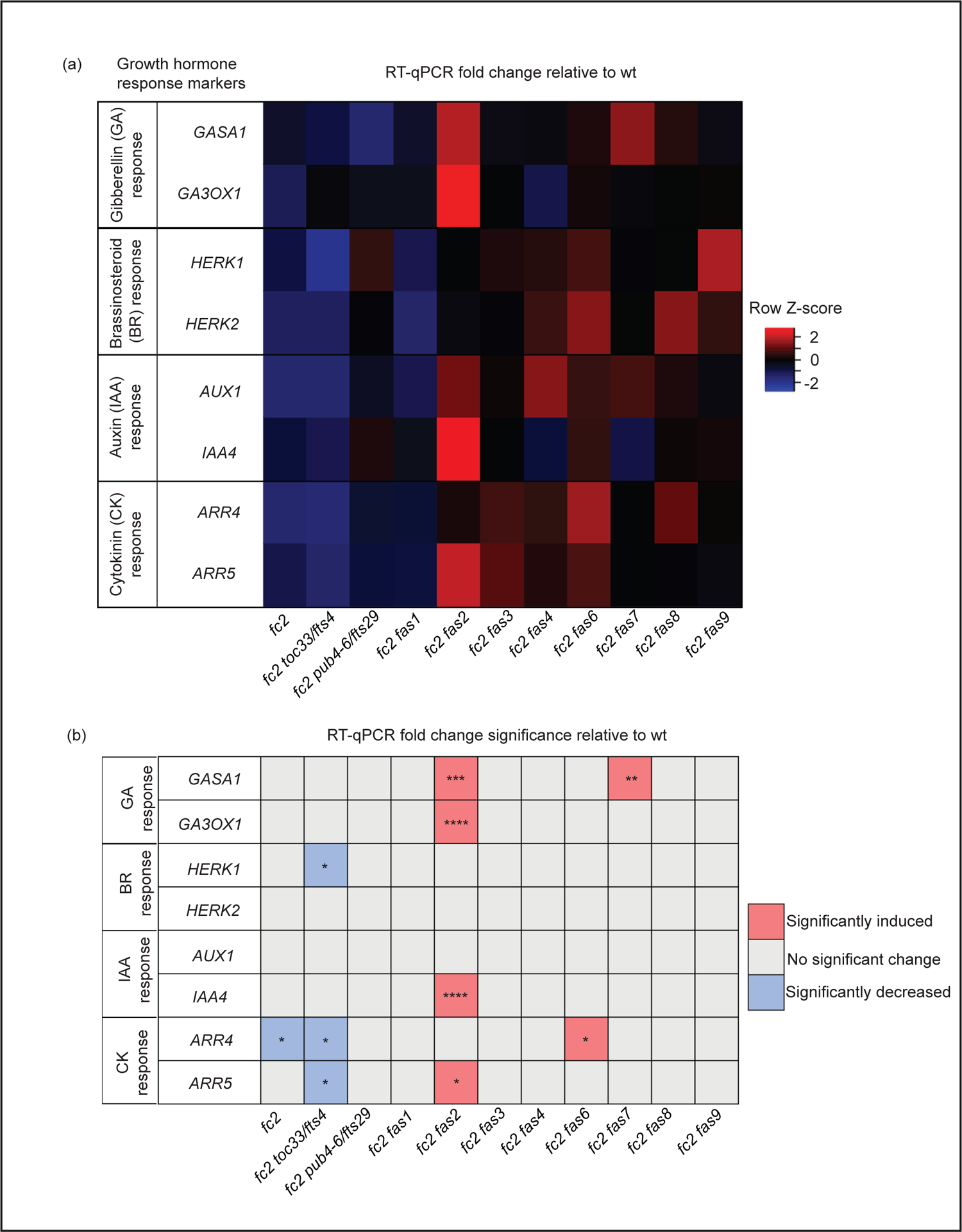
Growth hormone response markers are perturbed in some *fc2 fas* mutants Expression of growth hormone response marker genes was probed via RT-qPCR. Transcript abundance was monitored in 21-day-old plants grown for 14 days in constant light conditions and then transferred to cycling light conditions (16h light/8h dark) for 7 days. Gibberellin (GA), brassinosteroid (BR), auxin (IAA), and cytokinin (CK) response marker genes (two of each) were selected from the literature (see main text) to probe growth hormone response. (a) A heatmap table summarizing growth hormone response fold-change relative to wt. Shades of red indicate increased transcript abundance; shades of blue indicate decreased transcript abundance. All quantification of RT-qPCR analyses were tested using a one-way ANOVA. A Dunnett’s multiple comparisons post hoc were used to compare variation between genotypes relative to wt (* = *P* ≤ 0.05, ** = *P* ≤ 0.01, *** = *P* ≤ 0.001, **** = *P* ≤ 0.0001). n = 3 biological replicates. Error bars = +/- SEM. Closed circles indicate individual data points.

To investigate BR signaling, we chose probes for genes reported to respond to BR: *HERCULES RECEPTOR KINASE 1* (*HERK1*) and *HERCULES RECEPTOR KINASE 1* (*HERK2*) (Guo et al., 2009). Here, we did not observe any significant changes of expression between wt and *fc2*, or any *fc2 fas* mutants relative to wt (**Figs. 5a and b**). Of the mutants tested, only *fc2 toc33* exhibited a significant decrease in the expression of *HERK1*.

To investigate IAA signaling, we chose probes for genes reported to respond to IAA: *AUXIN RESISTANT 1* (*AUX1*) and *INDOLE-3-ACETIC ACID INDUCIBLE 5* (*IAA5*) (Paponov et al., 2008). We did not observe a change in the expression of these two genes in *fc2* relative to wt (**Figs. 5a and b**). However, only *IAA5* was significantly induced in *fc2 fas2,* relative to wt.

Next, we chose probes for genes reported to respond to CK signaling: *ARABIDOPSIS RESPONSE REGULATOR 4* (*ARR4*) and *ARABIDOPSIS RESPONSE REGULATOR 5* (*ARR5*) (D’Agostino et al., 2000). Relative to wt, we observed repression of *ARR4* in *fc2* (**Figs. 5a and b**). This change was reversed in all *fc2 fas* mutants, except *fc2 toc33* and *fc2 fas6*. Relative to wt, *ARR5* was only significantly induced in *fc2 fas2*.

Together, these data suggest no obvious induction or repression pattern in GA, BR, IAA, and CK marker genes in *fc2* relative to wt. Furthermore, while the expression of these growth hormone marker genes is perturbed in a few *fc2 fas* mutants (*fc2 fas2*, *fc2 fas6*, *fc2 fas7*) relative to wt, there was no obvious pattern of differential expression of these genes amongst all the *fc2 fas* mutants. The one notable exception was *fc2 fas2*, which exhibited induction of GA, IAA, and CK response marker genes.

### Suppression of programmed cell death in fc2 fas mutants is not generally correlated with reduced growth

Previous studies demonstrate that tolerance to abiotic stress is linked to reduced growth rates. For example, reduced GA signaling can achieve abiotic stress tolerance at the expense of growth, leading to dwarf phenotypes (Colebrook et al., 2014). Indeed, compared to *fc2*, four *fc2 fas* mutants (*fc2 fas1, fc2 fas2*, *fc2 fas4*, and *fc2 fas7*) have smaller leaves with altered morphology (**Fig. 6a**) and fail to rescue the impaired growth phenotype of cycling light-stressed *fc2* (**Fig. 1d**). Furthermore, *fc2 fas2* and *fc2 fas7* have altered GA marker gene expression (**Figs. 5a and b**). Together, these observations open the possibility that the suppression of PCD in *fc2* can be achieved by reducing growth rates. To test this possibility, we applied exogenous GA_3_ treatment. Under constant light conditions, the GA_3_ treatment led to wt, *fc2*, and *fc2 fas1*, *fc2 fas2*, *fc2 fas7* mutants (but not *fc2 fas4*) having visually longer petioles than untreated plants (indicative of response to GA_3_) (**Fig. 6b**). The same lines showed a GA_3_-dependent increase in biomass (**Fig. 6c**). However, this difference was only statistically significant for *fc2 fas2*, suggesting that this mutant may be a GA-sensitive dwarf.

**Figure 6.**
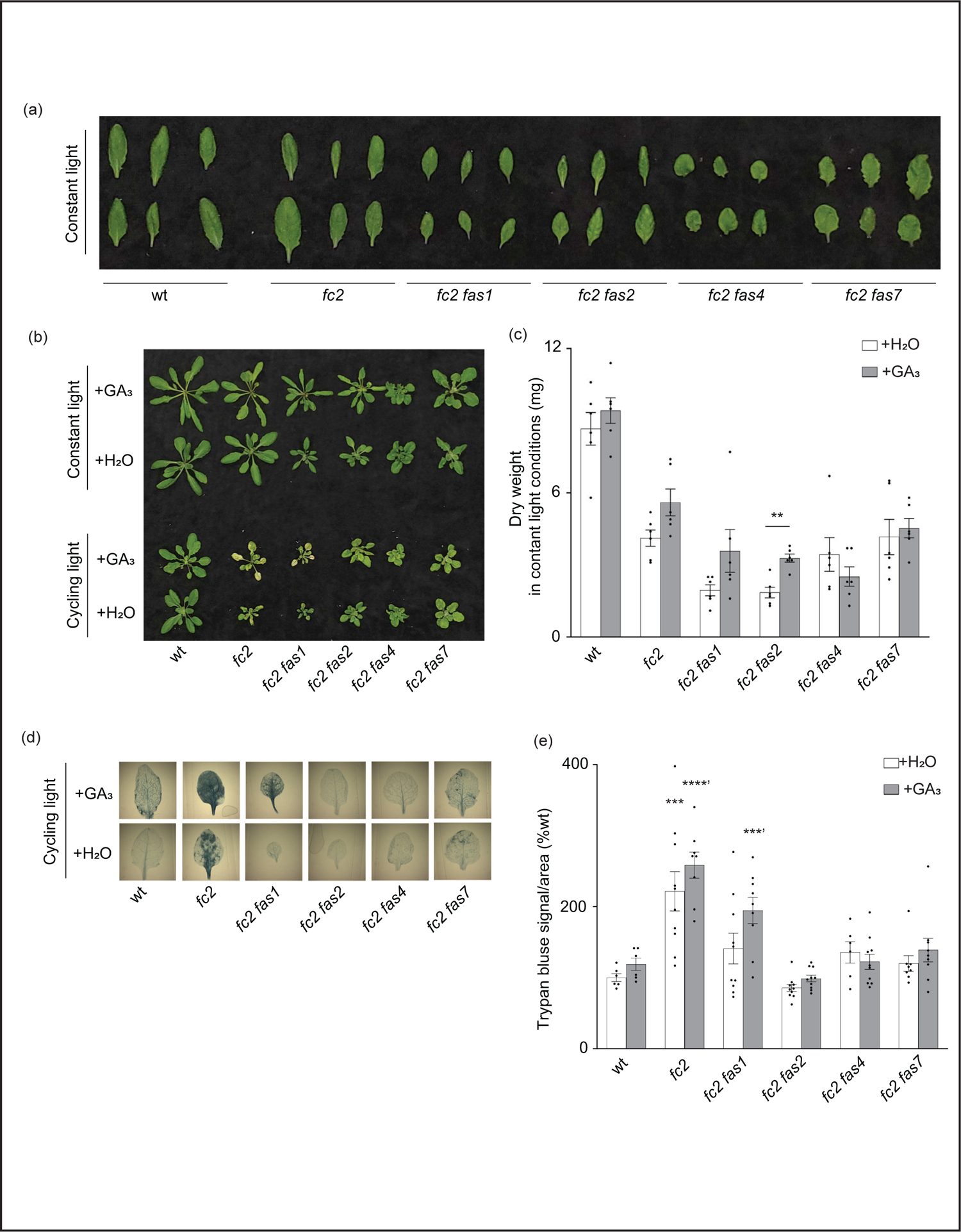
Response of *fc2 fas* mutants to exogenous GA_3_ treatments The ability of gibberellin (GA) to promote programmed cell death in select *fc2 fas* mutants (those with growth deficiencies) was tested. Plants were grown for 21 days in constant light conditions or 14 days in constant light conditions and then transferred to cycling light conditions (16h light/8h dark) for 7 days. Exogenous GA_3_ (10^-4^ M GA_3_) treatment was administered starting on day 7. (a) Individual leaves from 21-day-old plants grown in constant light conditions. (b) Representative images of plants grown in constant light conditions (top), or constant light conditions followed by cycling light conditions (bottom), with and without GA_3_ treatment. (c) Dry weight biomass (mg) was collected from plants grown in constant light, with H_2_O (mock control) or GA_3_ treatment. Student t-tests were used to compare differences between treatments (n = 6 biological replicates). (d) Representative images of leaves from plants grown under cycling light stress conditions, with H_2_O or GA_3_ treatment, that were stained with trypan blue. The dark blue color indicates cell death. (e) Quantification of trypan blue stains shown in d. Separate one-way ANOVAs and Dunnett’s multiple comparisons post hoc were used to compare variation between genotypes (relative to *fc2*) treated with H_2_O or with GA_3_ (** = *P* ≤ 0.01, *** = *P* ≤ 0.001, **** = *P* ≤ 0.0001). n ≥ 10 leaves from individual plants. Error bars = +/- SEM. Closed circles indicate individual data points.

Next, we explored the possibility that exogenous GA_3_ treatment can affect reduced PCD in the *fc2 fas* mutants. Under cycling light conditions, GA_3_ treatment restored PCD in the *fc2 fas1* mutant (**Figs. 6b, d, and e)**. However, the other *fc2 fas* mutants (*fc2 fas2*, *fc2 fas4*, and *fc2 fas7*) still retained their ability to suppress PCD when treated with GA_3_. These data suggest that the suppression of PCD observed in *fc2 fas2*, *fc2 fas4*, and *fc2 fas7* is not chemically coupled to GA_3_-regulated growth enhancement. In contrast, the suppression observed in *fc2 fas1* may be due, at least in part, to a GA deficiency and/or a reduced growth rate. Together, these results indicate that a reduced growth rate can protect from PCD in *fc2*, but it is not a general mechanism among the eight *fas* mutations.

### fas mutations perturb stress hormone signaling in the fc2 mutant

The stress phenotypes of the *fc2 fas* mutants suggest that stress hormone pathways may be altered. To test this, we focused on the transcriptional responses to four major plant stress hormones, Salicylic Acid (SA), Jasmonic Acid (JA), Abscisic Acid (ABA), and Ethylene (ET), which have all been shown to be regulated in response to ^1^O_2_ and play roles in PCD and senescence (Laloi and Havaux, 2015).

To assess SA responses, we chose to probe the SA response genes *PATHOGENESIS-RELATED GENE 1* (*PR1*), *PATHOGENESIS-RELATED GENE 2* (*PR2*), and *PATHOGENESIS-RELATED GENE 5* (*PR5*) (Schmitz et al., 2010). *PR1* and *PR5* (but not *PR2*) were induced in *fc2* relative to wt (**Figs. 7a and b**). This induction was mostly reversed in the *fc2 fts* and *fc2 fas* mutants, except for *PR5* still being induced, relative to wt, in *fc2 fas4* and *fc2 fas7*. However, while analyzing the induction of *PR2,* we noticed that *fc2 fas8* had a very large induction of this gene (>260-fold compared to wt). Excluding *fc2 fas8* from our analysis, *PR2* was determined to be significantly (*P* ≤ 0.01) induced in *fc2* but not in any other *fc2 fas* mutant compared to wt **(Table S5)**. These data suggest that SA signaling is induced in *fc2* and is generally returned to wt levels in *fc2 fas* mutants.

**Figure 7.**
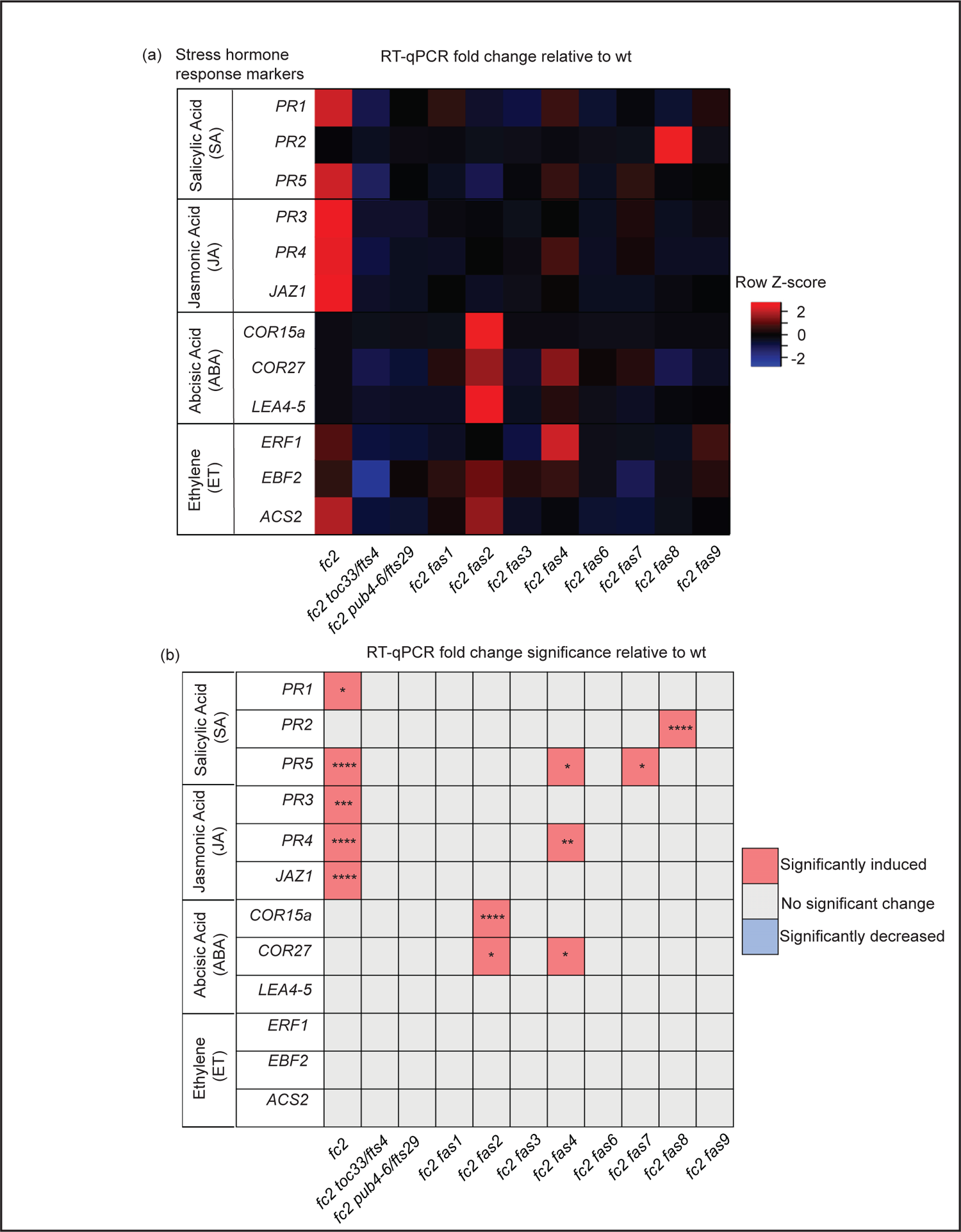
Analysis of stress hormone signaling in *fc2* and *fc2 fas* mutants Stress hormone signaling in *fc2* and *fc2 fas* mutants was assessed by measuring hormone response marker gene expression by RT-qPCR. Steady-state transcript levels were monitored in plants grown for 14 days in constant light conditions and then transferred to cycling light (16h light/8h dark) conditions for 7 days. Marker genes for salicylic acid (SA), Jasmonic acid (JA), abscisic acid (ABA), and ethylene (ET) were selected to probe stress hormone responses. (a) A heatmap table summarizing stress hormone response transcript abundance fold change relative to wt. Shades of red indicate increased transcript abundance and shades of blue indicate decreased transcript abundance. (b) Table reporting the significance of the difference in stress hormone response marker transcript abundance relative to wt. All quantification of qPCR analyses were tested using a one-way ANOVA. A Dunnett’s multiple comparisons post hoc were used to compare variation between genotypes relative to wt (* = *P* ≤ 0.05, ** = *P* ≤ 0.01, *** = *P* ≤ 0.001, **** = *P* ≤ 0.0001). n = 3 biological replicates.

To assess transcriptional responses to JA, we chose to monitor marker genes *PATHOGENESIS-RELATED GENE 3* (*PR3*), *PATHOGENESIS-RELATED GENE 4* (*PR4*), and *JASMONATE-ZIM-DOMAIN PROTEIN 1* (*JAZ1*) (Schmitz et al., 2010, Valenzuela et al., 2016). All three genes were significantly induced in *fc2*, relative to wt (**Figs. 7a and b**). This induction pattern was reversed in *fc2 toc33*, *fc2 pub4-6*, and all *fas* mutants, except for *PR4* in *fc2 fas4*. As such, JA signaling may be activated in *fc2* but is reversed by the *fts* and *fas* mutations.

To assess transcriptional responses to ABA, we probed the marker genes *COLD-REGULATED 15A* (*COR15A*), *COLD REGULATED GENE 27* (*COR27*), and *LATE EMBRYOGENESIS ABUNDANT 4-5* (*LEA4-5*) (de Torres-Zabala et al., 2007, Hundertmark and Hincha, 2008, Hoth et al., 2002). We observed no significant induction of these ABA markers in *fc2* relative to wt (**Figs. 7a and b**). The only marker genes induced, relative to wt, in *fc2 fas* mutants were *COR15a* in *fc2 fas2* and *COR27* in *fc2 fas2* and *fc2 fas4*. These data suggest that ABA signaling is not activated in *fc2*, but may be altered in *fc2 fas2* and *fc2 fas4*.

Finally, to assess transcriptional responses to ET, we probed the ET signaling marker genes *ETHYLENE RESPONSE FACTOR 1* (*ERF1*), *EIN3-BINDING F BOX PROTEIN 2* (*EBF2*), and *1-AMINO-CYCLOPROPANE-1-CARBOXYLATE SYNTHASE 2* (*ACS2*) (Muller and Munne-Bosch, 2015, Yu et al., 2021). We did not observe any induction of these ET response genes in any mutant relative to wt, suggesting that ethylene signaling is not broadly affected in *fc2* or perturbed by the *fas* mutations (**Figs. 7a and 7b**).

As SA and JA hormone response markers can play a role in senescence (Morris et al., 2000, He et al., 2002), we also chose to investigate the expression of known senescence-associated genes (SAGs) (Niu et al., 2020). Here, *SAG12*, *SAG13*, *SAG21*, *SAG101*, and *WRKY53* were probed by RT-qPCR. Only *SAG13* and *SAG21* were induced in *fc2* relative to wt **(Figs. S5a and b)**. The expression of *SAG13* and *SAG21* was reduced to levels statistically insignificant from wt in all fc2 suppressor mutants tested. *SAG101,* on the other hand, was observed to be induced in *fc2 fas4, fc2 fas7*, and *fc2 fas9* relative to wt. Together, these results suggest that SA and JA signaling is induced in stressed *fc2* and returned to wt levels in most *fc2 fts* and *fc2 fas* mutants. Thus, the stress response in *fc2* may also involve the activation of an early senescence program.

#### *Testing the tolerance of* fc2 fas *mutants to additional forms of oxidative stress*

As *fas* mutants suppress ^1^O_2_ signaling phenotypes, we next asked if this suppression is specific to ^1^O_2_ overproduction in the chloroplast or if any of these mutants are generally tolerant to photooxidative stress. To investigate this possibility, we first tested the response of *fas* mutants to two forms of abiotic stress that are expected to produce different forms of ROS in the chloroplast: EL and methyl viologen (MV). EL treatment produces high levels of ^1^O_2_ at PSII (Vass et al., 1992), while MV treatment produces superoxide (O_2_^-^) at PSI (Hassan, 1984), followed by the spontaneous or enzymatic conversion of O_2_^-^ to H_2_O_2_ (Hassan, 1988).

To test the tolerance of *fas* mutants to EL, 21-day-old plants were subjected to 1450-1500 µmol photons s^-1^ m^-2^ for 24 hours. As EL can generate heat, we set the chamber to 10°C to offset this effect (the resulting average leaf temperature was 19.7°C). Here, both wt and *fc2* exhibited photobleaching and lesion formation in leaf tissue, which failed to recover 3 days after 24h EL treatment (**Fig. 8a**). In contrast, *fc2 pub4-6, fc2 fas1*, *fc2 fas2*, *fc2 fas7*, and *fc2 fas8* exhibited delayed lesion formation. To quantify this response, lesion count ratios (# damaged leaves/# total leaves) were calculated from plants immediately after 24 hours of EL (prior to recovery to limit potential regreening and/or new growth). We observed no significant difference in the lesion count ratio between wt and *fc2*, both having an average lesion count ratio of ∼0.54 (**Fig. 8b**). The formation of necrotic lesions was observed to be delayed (relative to *fc2*) in *fc2 pub4-6* and *fc2 fas1*, *fc2 fas2*, *fc2 fas7*, and *fc2 fas8*. While conducting lesion counts, we observed that leaf lesions were not always uniform between different genotypes (e.g., in *fc2 fas3* and *fc2 fas6*, lesions appeared larger on the leaves of some plants (**Fig. 8a**). To quantify further EL-induced cell death, trypan blue stains were performed. Again, no significant difference was observed between wt and *fc2* (**Figs. 8c and d**). We did observe a significant delay in cell death (relative *fc2*) in *fc2 pub4-6, fc2 fas1*, *fc2 fas2*, *fc2 fas7*, *fc2 fas8,* and *fc2 fas9*, but not in *fc2 toc33*, *fc2 fas3*, and *fc2 fas6*.

**Figure 8.**
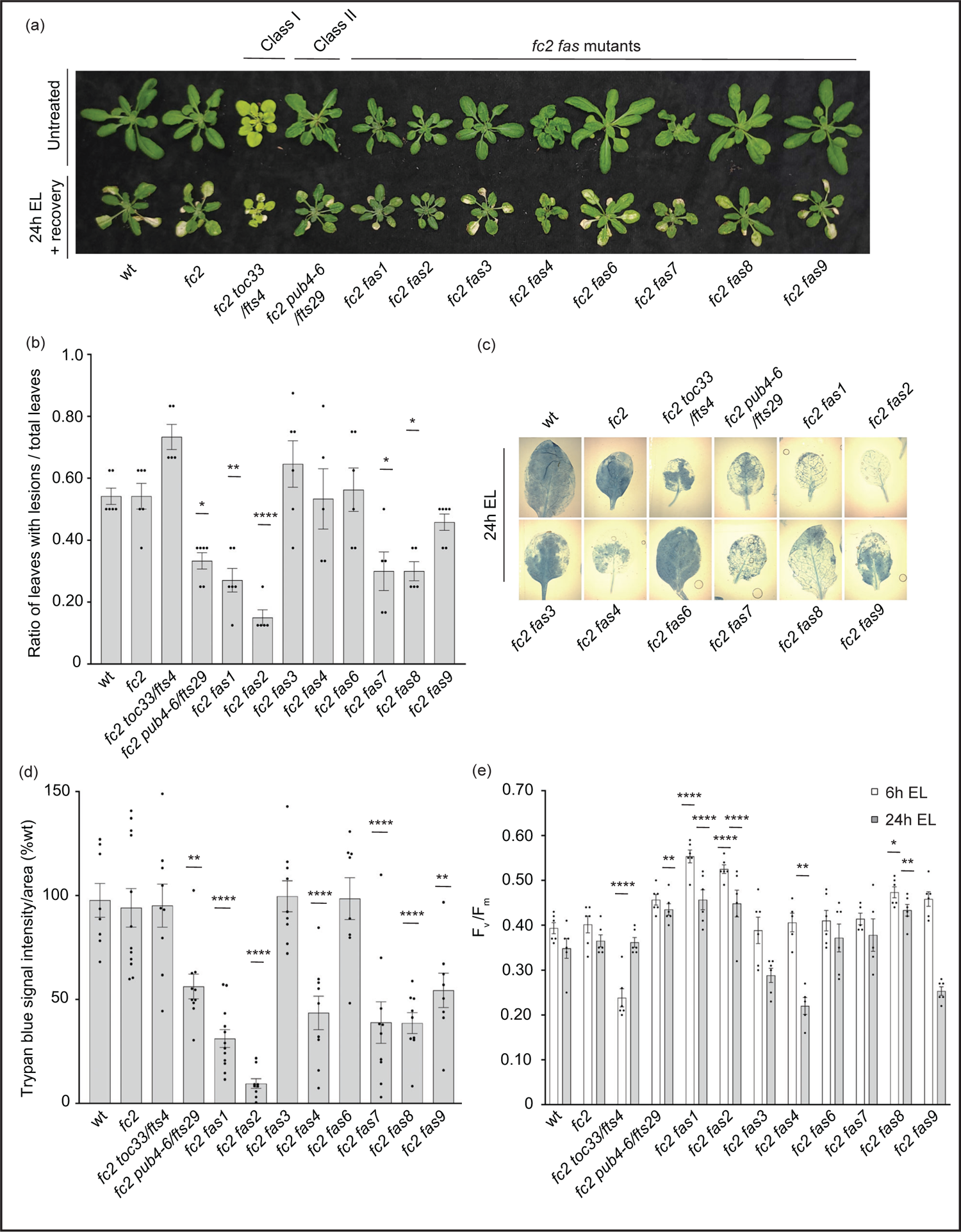
Testing the tolerance of *fc2 fas* mutants to excess light stress Excess light (EL) treatments were applied to test the tolerance of *fc2 fas* mutants to a different source of chloroplast reactive oxygen species (ROS). Plants were grown for 21 days in constant light conditions and then exposed to EL at an intensity of 1450-1550 µmol photons m^-2^ sec^-1^ white light at 10 °C. (a) Image showing representative 25-day-old plants, either unexposed (top row) or exposed to EL stress for 24 hours and allowed to recover for three days (bottom row). (b) Quantification of lesion counts (ratio of leaves with observable cell death/ healthy leaves) immediately after 24h EL exposure, prior to regreening and new growth (n ≥ 5 replicate plants). Prior to treatment, plants had no observable lesions. (c) Representative images of leaves collected (from plants assessed for lesion counts in panel b) and stained with trypan blue. The dark blue color indicates dead tissue. (d) Quantification of trypan blue stains shown in c (n ≥ 10 leaves from replicate plants). (e) Quantification of maximum photosynthetic efficiency (F_v_/F_m_) after 6h or 24h EL exposure (n ≥ 5 replicate plants). Prior to treatment, plants had the expected F_v_/F_m_ ratios for unstressed plants (0.80-0.84) (**Fig. S6a**). All F_v_/F_m_ measurements, lesion counts, and trypan blue stains were tested with a one-way ANOVA and Dunnett’s multiple comparisons post hoc to compare variation between genotypes relative to *fc2* (* = *P* ≤ 0.05, ** = *P* ≤ 0.01, **** = *P* ≤ 0.0001). Error bars = +/- SEM. Closed circles indicate individual data points.

We also measured the impact of EL stress on photosynthesis by calculating the maximum potential quantum efficiency of PSII (F_v_/F_m_). Generally, decreased F_v_/F_m_ ratios correlate with plant stress. To set baseline F_v_/F_m_ values for unstressed plants and to ensure that *fas* mutants are not starting in a pre-stressed state, chlorophyll fluorescence measurements were taken from untreated, constant light grown plants at 21, 23, and 28 days old (21-28 days being the general age of plants assessed via chlorophyll fluorescence measurements in this study). The F_v_/F_m_ values ranged from ∼0.81 to 0.84 **(Fig. S1a-c).** The only mutant with constitutively decreased values compared to *fc2* was *fc2 toc33*, which ranged between ∼0.78-0.81.

We next measured F_v_/F_m_ values of 21-day-old plants immediately after 6h and 24h EL treatment. After 6h EL treatment, F_v_/F_m_ values decreased in all plant lines (**Fig. 8e**). However, we observed no significant difference between *fc2* and wt. Compared to *fc2*, we did observe significantly lower F_v_/F_m_ values in *fc2 toc33* and significantly higher F_v_/F_m_ values in *fas1*, *fc2 fas2*, and *fc2 fas8*. Following 24h EL treatment, F_v_/F_m_ values decreased even further in most plant lines. Again, we observed no significant difference between *fc2* and wt. However, at this time point, compared to *fc2*, we observed significantly lower F_v_/F_m_ values in *fc2 fas4* and significantly higher F_v_/F_m_ values in *fc2 pub4-6*, supporting earlier observations that the *pub4-6* mutation protects cells from EL stress (Tano et al., 2023). *fc2 fas1*, *fcs fas2*, and *fc2 fas8* also had significantly higher F_v_/F_m_ values compared to *fc2*. Together, these results show that six of the eight *fas* mutations also lead to a similar degree of photoprotection from EL.

To test the tolerance of *fas* mutants to a different form of photooxidative chloroplast stress, we next applied 20 μM or 200 μM MV to 21-day-old plants grown in constant light to induce the accumulation of O_2_^-^, and consequently H_2_O_2_ in chloroplasts. Lesion count ratios were calculated four days after treatment (to allow time for lesion formation) to assess MV tolerance. When 20 μM MV was applied, wt and *fc2* exhibited similar lesion formation levels (**Figs. 9a and b**). Only *fc2 fas2* and *fc2 toc33* had reduced lesion formation relative to *fc2*. None of the plants survived a more stringent (200 uM) MV treatment (**Fig. 9a**). As such, we did not assess lesion formation in response to this treatment regimen. When photoinhibition was measured via chlorophyll fluorescence 24 hours after 20 μM MV treatment, all plant lines exhibited a decrease in F_v_/F_m_ values (**Fig. 9c**). However, no significant difference was observed between *fc2* and wt. Compared to these controls, only *fc2 fas2* displayed a tolerance to 20 μM MV (**Fig. 9c**) (we did not assess photoinhibition in response to 200 μM MV). Together, these data suggest that *fc2 fas2*, but not other *fas* mutants, may have a general tolerance to chloroplast photooxidative stress.

**Figure 9.**
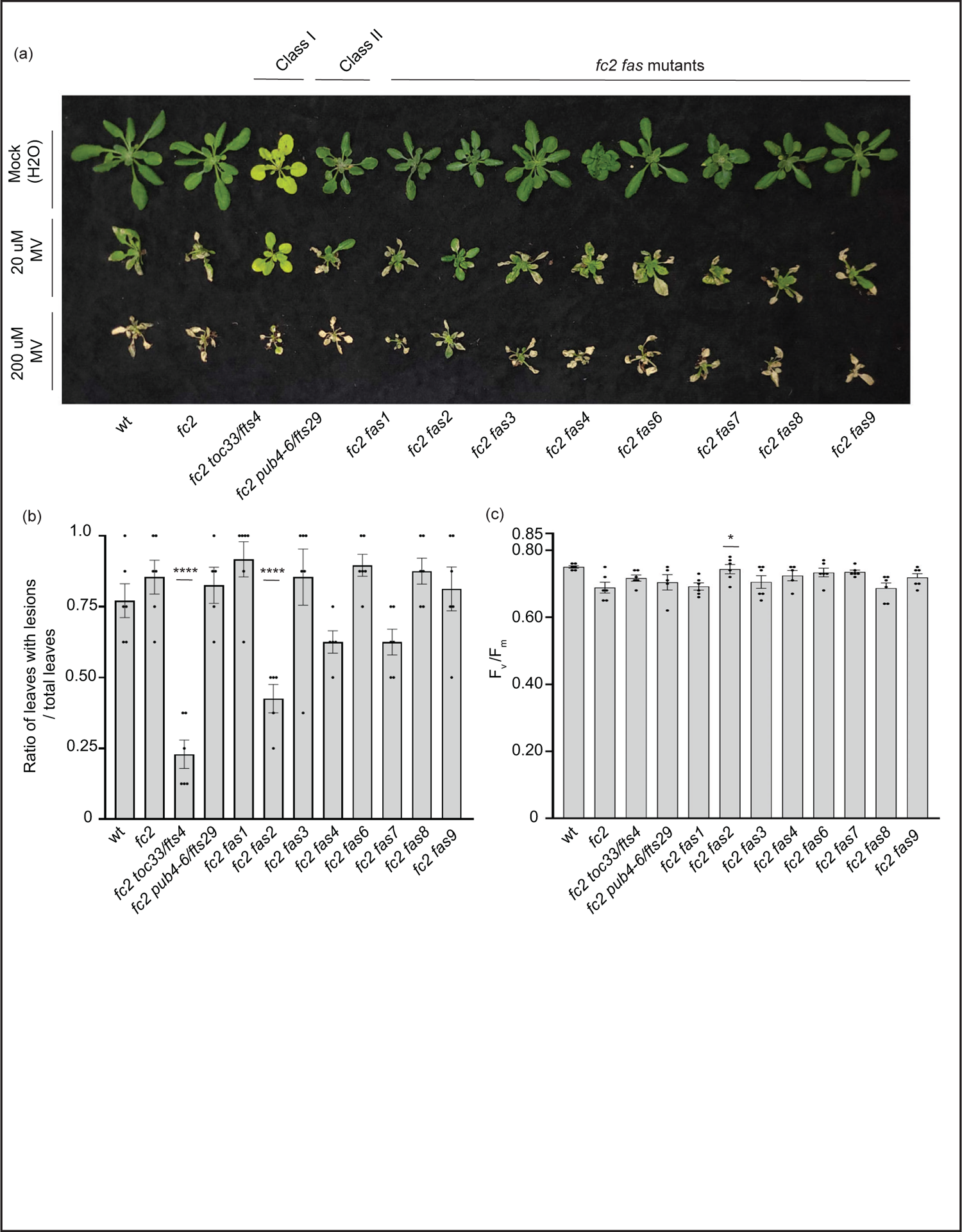
Testing the tolerance of *fc2 fas* mutants to methyl viologen Methyl viologen (MV) treatments were applied to test the tolerance of *fc2 fas* mutants to chloroplast hydrogen peroxide (H_2_O_2_). Plants were grown for 21 days in constant light conditions and then treated with MV to generate superoxide (^-^O_2_) and, subsequently, H_2_O_2_ stress. (a) Image of representative 25-day-old plants treated with either Mock (H_2_O) treated (top row), 20µm MV (middle row), or 200µm MV (bottom row) at 21 days and then grown for another four days in constant light conditions. (b) Quantification of lesion counts (ratio of leaves with observable cell death/ healthy leaves) from plants in panel a (n ≥ 4 plants). (c) Quantification of maximum photosynthetic efficiency (F_v_/F_m_) 24h hours after MV exposure (n ≥ 4 plants). Prior to treatment, plants had the expected F_v_/F_m_ ratios for unstressed plants (∼0.80-0.84) (**Fig. S6a**). All lesion count and F_v_/F_m_ measurements were tested with a one-way ANOVA and Dunnett’s multiple comparisons post hoc to compare variation between genotypes relative to *fc2* (n.s. = not significant, * = *P* ≤ 0.05, **** = *P* ≤ 0.0001). Error bars = +/- SEM. Closed circles indicate individual data points.

#### *Testing the tolerance of* fc2 fas *mutants to heat, freezing, and carbon starvation stress*

To test if *fas* mutations are specifically affecting responses to chloroplast ^1^O_2_ stress or if they are generally tolerant to abiotic stress, we next tested their response to heat (40°C), freezing (−20°C), and dark-induced carbon starvation (for 5-7 days). Such abiotic stress can negatively affect chloroplast function and enhance light-dependent accumulation of chloroplast ROS (Gururani et al., 2015). As such, all three stresses were tested in dark conditions to limit the possible role of light-dependent chloroplast ROS.

For heat treatments, 21-day-old plants were exposed to 40°C in the dark for 16 or 24 hours, and then plants were allowed to recover for three days to assess survival. Plant survival was variable after 16 hours at 40°C. After 24h at 40°C, however, wt and *fc2* plants were unable to recover (**Fig. 10a**). While most *fc2* suppressors also failed to recover, *fc2 pub4-6* and *fc2 fas2* were both able to survive. To quantify the level of stress incurred by the plants, we again measured F_v_/F_m_ values, which were taken 1 hour after the heat treatment. After 16h of heat, all lines showed a decrease in F_v_/F_m_ values, indicating a degree of photosynthetic stress (**Fig. S7a**). However, we observed no significant difference between wt and *fc2*. Most *fc2 fts*/*fas* mutants had similarly low F_v_/F_m_ values. However, *fc2 pub4-6* and *fc2 fas2* showed a degree of tolerance with significantly higher F_v_/F_m_ values (compared to *fc2*). Interestingly, the *fcs toc33* mutant appeared sensitive to the heat treatment and had significantly lower F_v_/F_m_ values than *fc2*. When exposed to a longer heat treatment of 40°C for 24 hours, most of the plant lines still experienced a pronounced reduction of F_v_/F_m_ values (**Fig. 10b**). Again, *fc2 pub4-6* and *fc2 fas2* still exhibited a significantly higher F_v_/F_m_ values than *fc2*, while *fc2 toc33* exhibited a significantly lower F_v_/F_m_ values than *fc2*. Together, these data suggest that the abiotic stress tolerance of *fts fas* mutants does not generally extend to heat tolerance. However, *fc2 fas2* and *fc2 pub4-6* can protect their photosynthetic machinery during prolonged heat stress.

**Figure 10.**
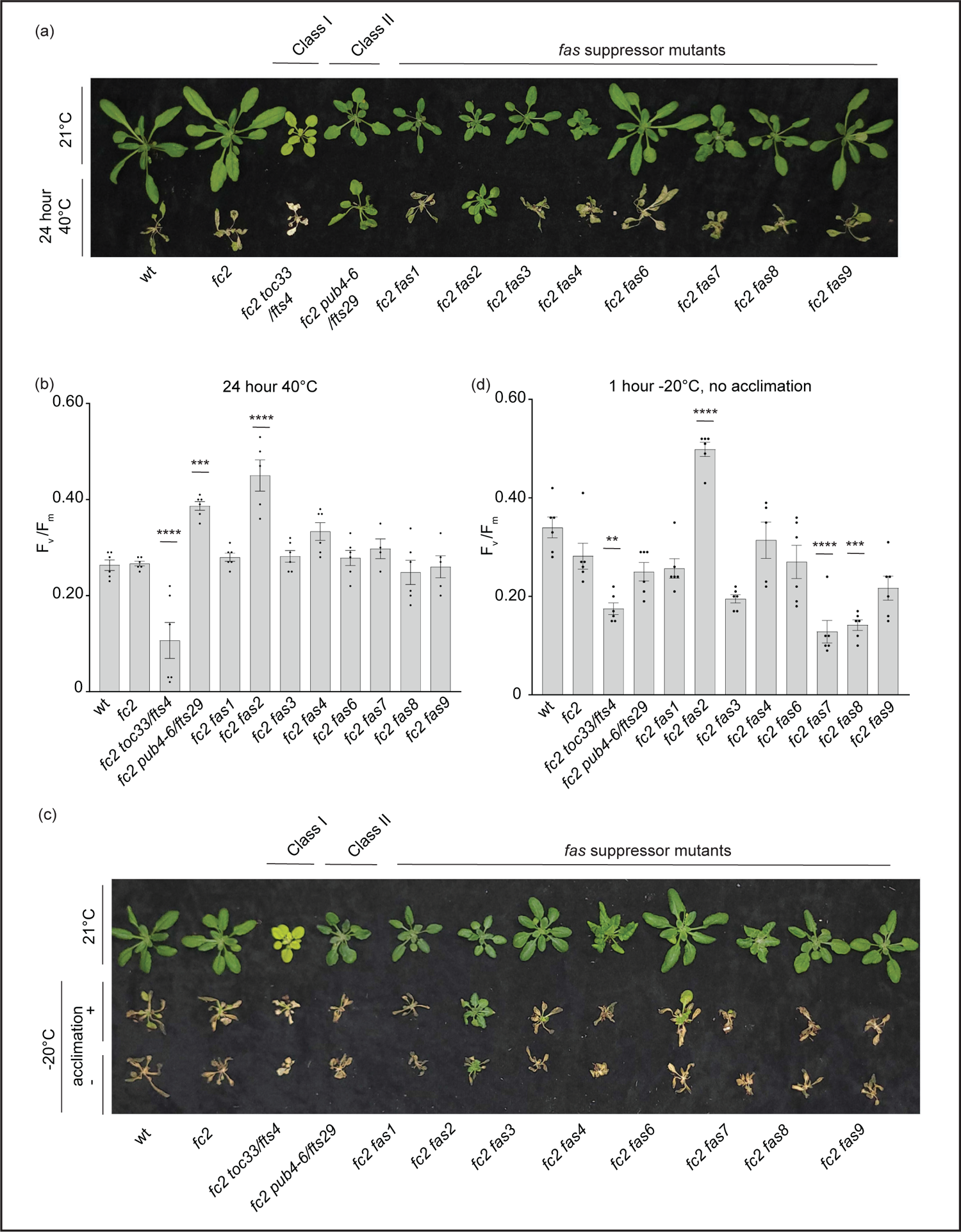
Testing the tolerance of *fc2 fas* mutants to heat and freezing stresses *fc2 fas* mutants were tested for their tolerance to different temperature stresses. Plants were grown for 21 days in constant light conditions and then exposed to heat stress (40°C) (a-b) or freezing stress (−20°C) (c-d). (a) Image showing representative 25-day-old plants, either unexposed (top) or exposed to 40°C for 24h and allowed to recover in constant light at 21°C for three days in constant light conditions. (b) Quantification of maximum photosynthetic efficiency (F_v_/F_m_) after 24h of 40°C treatment and one hour of recovery (n ≥ 4 plants). (c) Images of 25-day-old plants untreated (top row), exposed to freezing (−20°C for 1 hour) with cold acclimation for 16h at 4°C (middle row), or exposed to freezing without cold acclimation (bottom row) and allowed to recover in constant light at 21°C for three days in constant light conditions. (d) Quantification of F_v_/F_m_ measurements taken after freezing treatment (without prior acclimation) and allowed to recover for 2 hours (n ≥ 4 plants). All F_v_/F_m_ measurements were tested with a one-way ANOVA and Dunnett’s multiple comparisons post hoc to compare variation between genotypes relative to *fc2* (** = *P* ≤ 0.01, *** = *P* ≤ 0.001, **** = *P* ≤ 0.0001). Error bars = +/- SEM. Closed circles indicate individual data points.

To test mutants for tolerance to freezing stress, 21-day-old plants were first acclimated to cold (16h at 4°C) and then subjected to −20°C for 1 hour in the dark. When allowed to recover for 3 days, most plants failed to survive (**Fig. 10c**). However, *fc2 fas2* plants consistently exhibited a degree of survival and remained green. To quantify this stress response, we measured F_v_/F_m_ values of plants 1 hour after freezing treatment. As expected, all plants showed a decrease in F_v_/F_m_ values, but they were significantly higher in *fc2 fas2* compared to wt or *fc2* (**Fig. S7b**), suggesting that *fc2 fas2* mutants can protect their chloroplast membranes during freezing stress. We then increased the stringency of the experiment by exposing plants to the same freezing stress without prior cold acclimation. Surprisingly, *fc2 fas2* mutants again survived (retained some green leaves) and retained higher F_v_/F_m_ values than wt or *fc2* (**Figs. 10c and d**). Under these conditions, *fc2 toc33, fc2 fas7,* and *fc2 fas8* exhibited significantly decreased F_v_/F_m_ values. Together, these data suggest that the *fc2 fas* mutants are not generally tolerant to freezing stress (*fc2 fas7* and *fc2 fas8* are sensitive), but *fc2 fas2* continues to show a broad tolerance to multiple types of abiotic stress.

Dark-induced carbon starvation activates senescence, autophagy, and chlorophagy and predominantly involves H_2_O_2_ signaling (Wada et al., 2009, Perez-Perez et al., 2012). To test the response of *fas* mutants to dark-induced carbon starvation, 23- or 21-day-old plants were placed in the dark for five or seven days, respectively, and then allowed to recover for four days. We also included *autophagy 5* (*atg5*, a mutation that impairs the assembly of autophagosomes and blocks canonical autophagy pathways (Thompson et al., 2005)) in the wt (*atg5*) and *fc2* (*fc2 atg5*) backgrounds to control dark-induced carbon starvation sensitivity. After five days of dark starvation, wt and *fc2* plants accumulate leaf lesions but survive (retain green tissue) (**Fig. 11a**). Consistent with previous work (Thompson et al., 2005), the carbon starvation-sensitive mutant *atg5* (as well as the *fc2 atg5* double mutant) failed to survive after being subjected to only five days of darkness (no green tissue remained). The *fc2 fts* and *fc2 fas* mutants displayed different levels of survival. To quantify this, we measured F_v_/F_m_ values 1 hour after dark treatment. Both wt and *fc2* showed a reduction of F_v_/F_m_ values but were statistically similar. However, after five days of dark treatment, *fc2 toc33*, *fc2 fas2,* and *fc2 fas7* exhibited a higher F_v_/F_m_ than *fc2* (**Fig. 11b**). Under a more stringent regimen (7 days dark), all plants failed to survive, except *fas2 fc2* and *fas7 fc2* (**Fig. 11a**). Only *fc2 fas7* was observed to have a significantly increased F_v_/F_m_ values compared to *fc2* (**Fig. 11b**). Thus, while *fc2 toc33*, *fc2 fas2,* and *fc2 fas7* are tolerant to dark-induced carbon starvation and display a “stay-green” phenotype, the remaining six *fc2 fas* mutants are not tolerant to these stresses.

**Figure 11.**
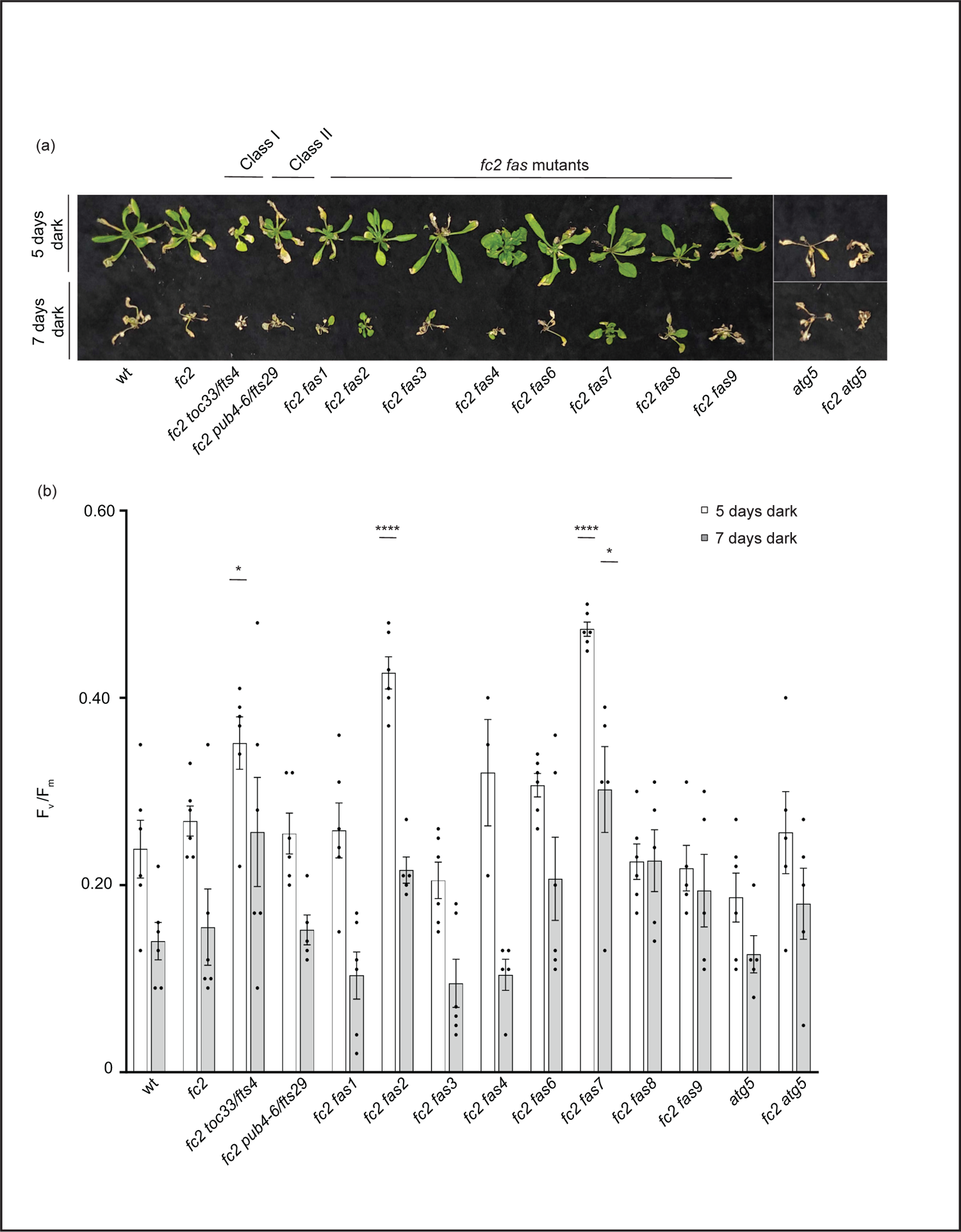
Testing the tolerance of *fc2 fas* mutants to dark-induced carbon starvation The tolerance of *fc2 fas* mutants to dark-induced carbon starvation was assessed. (a) Representative images of 32-day-old plants exposed to dark-induced carbon starvation for five (starting at 23 days old) or seven days (starting at 21 days old). Plants were allowed to recover for four days in constant light conditions. (b) Quantification of maximum photosynthetic efficiency (F_v_/F_m_) 1 hour after removal from dark conditions. All F_v_/F_m_ measurements from each treatment regimen were tested separately with a one-way ANOVA with Dunnett’s multiple comparisons post hoc to compare variation between genotypes relative to *fc2* (* = *P* ≤ 0.05, **** = *P* ≤ 0.0001). n ≥ 4 plants. Error bars = +/- SEM. Closed circles indicate individual data points.

Taken together, these data suggest that the *fc2 fas* mutants are only generally tolerant to abiotic stresses that produce ^1^O_2_ in chloroplasts (*fc2* stress and EL) and not to those that produce H_2_O_2_ in the chloroplast (MV) or ROS outside the chloroplast (heat, freezing, dark starvation). However, the *fc2 fas2* mutant proved to be an outlier, being tolerant to every stress we tested. A summary of these phenotypes is in **Table S9**.

## Discussion

The accumulation of chloroplast ^1^O_2_ can trigger retrograde signaling and cellular degradation, allowing a plant to respond to environmental stress by activating acclimation systems or promoting plant fitness by terminating dysfunctional cells (Wagner et al., 2004, Ramel et al., 2013, Woodson et al., 2015). The mechanisms behind these signals are not well understood, but the use of Arabidopsis genetic model systems such as *fc2*, which conditionally produce chloroplast ^1^O_2_, allow researchers to identify and characterize signaling components. To this end, a forward genetic suppressor screen using EMS had already yielded mutants that suppress chloroplast ^1^O_2_-signaling and PCD in *fc2* seedlings and has successfully identified proteins involved in ubiquitination (Woodson et al., 2015) and plastid gene expression (Alamdari et al., 2020, Alamdari et al., 2021) as being important in propagating the ^1^O_2_ signal from chloroplasts. However, these mutations were generated using EMS, which predominantly produces recessive alleles, and will mostly uncover positive regulators of ^1^O_2_-signaling. To “complement” this work and gain a more comprehensive view of the genes involved in chloroplast stress signaling, here we have used activation tagging to generate dominant gain-of-function alleles, which we hypothesized would identify new genes involved in ^1^O_2_-signaling, including negative regulators.

### Eight dominant fc2 fas mutants that suppress PCD reveal differential stage-specific ^1^O_2_ response mechanisms

The screen described above successfully identified eight dominant *fas* mutants that reduce ^1^O_2_-induced retrograde signaling and PCD in adult *fc2* plants. (**Figs. 1b and c**). However, only *fc2 fas3* and *fc2 fas7* block this PCD in the seedling stage (**Figs. 2a-c**), demonstrating that most of the *fas* mutations are stage-specific suppressors of PCD in *fc2*. This complication was not specific to the *fas* mutations. Here, we also demonstrated that some *fts* mutations also had a stage-specific effect: Class III mutants, which affect plastid gene expression and were isolated for their ability to suppress PCD in the seedlings stage, did not block PCD in the adult stage **(Fig. S4a-c).** ^1^O_2_ was previously shown to accumulate in *fc2 toc33* seedlings (Woodson et al., 2015). In adults, however, we observed ^1^O_2_ to accumulate (**Fig. 4a**), suggesting TOC33 may play different signaling roles in true leaves. Other examples of stage-specific blocking of ^1^O_2_-induced PCD have been reported. In *fc2* mutants, *ex1 executor 2* (*ex2*) blocked PCD and retrograde signaling in the seedling stage, but not in the adult stage. On the other hand, *oxi1* blocked PCD in the adult stage but not the seedling stage (Tano et al., 2023). The suppression of PCD by *ex1 ex2* appears to be indirect (reduced tetrapyrrole and ^1^O_2_ accumulation), but it is expected that OXI1 likely plays a signaling role in *fc2*. The mechanisms involved in these stage-specific differences are unknown, but previous studies have suggested that chloroplast biology and development may differ between cotyledon and true leaf mesophyll cells (Albrecht et al., 2006, Albrecht et al., 2008). Nonetheless, this study provides further evidence for the existence of multiple ^1^O_2_-signaling pathways that may be activated differentially, dependent on the age of the plant. This is intriguing, as it highlights the possibility that plants have evolved specific spatiotemporal strategies to trigger ^1^O_2_-induced PCD under multiple circumstances rather than a general ^1^O_2_-induced PCD pathway.

Our results showed that *fc2 fas* mutants display a wide range of visual phenotypes (**Fig. 1a**), stage-specific suppression of PCD (**Fig. 2a-d**), variation in growth- and stress-hormone responses (**Figs. 4a, 4b, 5a, and 5b),** and physiological responses to stress (**Figs. 8a-e, 9a-c, 10a-d, 11a-b)**. While the causative genes in each *fc2 fas* mutant have yet to be identified, these observations suggest that the activation tag influences different genes in each mutant. Identifying the causative genes should allow researchers to elucidate the molecular mechanisms by which ^1^O_2_ signals induce PCD, chloroplast turnover, and retrograde signaling.

### Singlet oxygen signaling is affected in fc2 fas mutants

An analysis of bulk ^1^O_2_ levels in the *fc2 fas* mutants showed that most still accumulate high levels of ^1^O_2_ in cycling light conditions (**Figs. 4a and b**). The exceptions are *fc2 fas2* and, to an extent, *fc2 fas4*. This suggests that most of the *fas* mutations suppress PCD by blocking a chloroplast signal and further supports that chloroplast ^1^O_2_-initiated PCD is a genetically encoded pathway in plants. We do not yet know the causal genes in the mutants, but they seem unlikely to be involved in tetrapyrrole accumulation (like class I *fts* mutants) or plastid gene expression (like class III *fts* mutants) as they show no obvious signs of reduced chloroplast development as seedlings or adults (e.g., pale phenotypes and reduced chlorophyll accumulation). Thus, it may be expected that the *fas* mutations act downstream and/or outside the chloroplast. The dominant nature of these alleles further suggests a negative signaling role for affected gene products. Interestingly, except for *fas4*, these mutations also block most of the retrograde signal to the nucleus (**Figs. 3a and b**), further indicating that ^1^O_2_-induced chloroplast retrograde signaling and PCD are coupled. The exact relationship between retrograde signaling and PCD is unknown in any system (Woodson, 2022), and the identification of the causative mutations in these mutants will shed light on the signaling mechanisms involved.

### ^1^O_2_ stress tolerance is not coupled with reduced plant growth

Plant stress tolerance can be influenced by the amount of energy available to mount a response, and plants balance these energy stores by regulating growth hormone pathways controlled by GAs, BRs, IAAs, and CKs (Huot et al., 2014, Zhang et al., 2020). None of our selected growth hormone response marker genes showed a pattern of induction or repression in stressed *fc2*. However, we did observe perturbation of GA-response genes in *fc2 fas2* and *fc2 fas7* (**Figs. 5a and b**). These two mutants (along with *fc2 fas1* and *fc2 fas4*) also did not reverse the impaired growth phenotype of cycling light-stressed *fc2* (**Fig. 1d**) and had smaller leaves with altered morphologies (**Fig. 6a**). As reduced levels of GA have been documented to lead to stress tolerance (Colebrook et al., 2014), we tested if the PCD suppression is due to reduced GA levels in these *fc2 fas* mutants. GA_3_ treatments revealed that PCD suppression in *fc2 fas1* is, at least partially, coupled with a reduced growth rate or a reduction in GA-signaling (**Figs. 6b-e**). Conversely, the PCD suppression in *fc2 fas2*, *fc2 fas4*, and *fc2 fas7* is not coupled with GA-signaling. Furthermore, *fc2 fas2* appears to be a classic GA-sensitive dwarf mutant, as its reduced growth can also be rescued via GA_3_ treatment. Together, these observations suggest that a general decrease in growth is not the primary mechanism of ^1^O_2_-induced PCD suppression in the *fc2 fas* mutants but, under some circumstances, may be able to increase tolerance to chloroplast ^1^O_2_ levels.

### ^1^O_2_-induced JA and SA signaling are generally mediated in fc2 fas mutants

A large body of evidence suggests SA and JA signaling is influenced by ^1^O_2_ production (D’Alessandro et al., 2020). For example, ^1^O_2_ accumulation in *flu* and *ch1* has been shown to enhance the accumulation of SA and JA, which have been hypothesized to affect ^1^O_2_-induced PCD. In *flu* mutants, a biotrophic defense protein that plays a role in SA accumulation, *EDS1* (*ENHANCED DISEASE SUSCEPTIBILITY 1*), is induced by ^1^O_2_, and PCD is suppressed in *flu eds1* double mutants (Ochsenbein et al., 2006). Reducing SA levels in *flu* also attenuates ^1^O_2_-induced PCD (Danon et al., 2005), further implicating a role for SA signaling as a positive regulator of ^1^O_2_-induced PCD. Blocking JA signaling in *flu* has led to conflicting results, with some studies suggesting JA may promote (Danon et al., 2005) or block (Przybyla et al., 2008) PCD in response to ^1^O_2_ accumulation in the chloroplast. In the *ch1* mutant, JA levels, but not SA levels, correlate to ^1^O_2_-induced PCD; the reduction of JA synthesis with the *dde2-2* mutation blocked EL-induced lesion formation in the *ch1* mutant (Ramel et al., 2013). Together, these studies show that there is likely a relationship between ^1^O_2_ and signaling by JA and SA, but it is complex.

For these reasons, we chose to investigate the role of stress hormone signaling in *fc2* and *fc2 fas* mutants. All tested JA-response markers probed were induced in stressed *fc2*, and this induction was reduced to wt levels in the *fc2* suppressors with only one exception (*PR4* in *fc2 fas4*) (**Figs. 7a and b**). A similar pattern was observed with SA marker gene expression. An interesting exception is the *fc2 fas8* mutant, which had a strong induction of *PR2*, suggesting that it may be playing a positive role in regulating SA and/or *PR2*. Together, these data are compelling as SA and JA signaling have documented roles in PCD and senescence (Morris et al., 2000, Hu et al., 2017). Indeed, some SAGs (*SAG13* and *SAG21*) are also induced in *fc2,* and this induction is reduced in most *fc2 fts/fas* mutants **(Fig. S8)**.

Together, these observations demonstrate that JA and SA stress hormone signaling often correlates to the amount of PCD in the *fc2* mutant when exposed to cycling light stress. It is unclear, however, if JA and SA induction is the cause or effect of cellular degradation pathways in *fc2*. Alternatively, SA and JA may be directly influenced by ^1^O_2_-induced retrograde signaling but then act in a parallel, but separate, pathway to PCD. Further studies into the role of SA and JA in *fc2* will help us better understand their roles in chloroplast ^1^O_2_-induced signaling, CQC, and PCD.

### Tolerance of fc2 fas mutants to chloroplast ROS, heat, freezing, and carbon-starvation stresses

Our genetic screen was designed to identify mutations that specifically alter chloroplast ^1^O_2_ stress signaling. However, we expect also to uncover mutations that led to a broad tolerance to photo-oxidative or abiotic stress and the multiple types of ROS that can be produced in and outside the chloroplast. To determine if this was the case, we tested the tolerance of *fc2 fas* mutations to EL and MV treatments, which leads to light-dependent ^1^O_2_ and H_2_O_2_ accumulation in the chloroplast, respectively (Triantaphylidès et al., 2008, Hassan, 1984) The responses of *fc2 fas* mutants to EL were complex. However, we did observe that *fc2 pub4-6* and nearly all *fas* mutations (excepting *fc2 fas3* and *fc2 fas6*) delay EL-induced photo-inhibition and/or PCD (**Figs. 8c and d**). This may indicate that EL is a natural stress that mimics what occurs in *fc2* mutants under cycling light conditions. However, the observation that *fc2 fas3* and *fc2 fas6* are not tolerant to EL suggests there are some differences we do not yet understand. In contrast to EL treatment, none of the *fc2 fas* mutants, except *fc2 fas2*, exhibited any tolerance to MV (**Figs. 9a-c**), suggesting *fas* mutations do not generally offer tolerance to chloroplast O_2_^-^ or H_2_O_2_. This may be expected as it does not appear that *fc2* mutants produce significant levels of H_2_O_2_ as seedlings (Alamdari et al., 2020). Furthermore, it has previously been shown that chloroplast H_2_O_2_ likely induces a retrograde signal separate from ^1^O_2_ (op den Camp et al., 2003).

To test for general stress tolerance, we chose three forms of abiotic stress: heat, freezing, and carbon starvation. Heat stress and freezing stress primarily involve H_2_O_2_ accumulation (Devireddy et al., 2021, Sachdev et al., 2021). Darkness-induced carbon starvation also involves H_2_O_2_ accumulation and activates senescence, autophagy, and chlorophagy (Wada et al., 2009, Perez-Perez et al., 2012, Izumi and Nakamura, 2017). When these additional stresses were tested, we observed that most *fc2 fts*/*fas* mutants behaved like wt and *fc2*, but a few interesting exceptions were noted **(Table S9)**. The *fc2 pub4-6* mutant was tolerant to heat stress, while *fc2 toc33* and *fc2 fas1* were observed to be sensitive to heat (**Figs. 10a and b**). The *fc2 fas7* mutant was also tolerant to carbon starvation and exhibited a “stay-green” phenotype (**Figs. 11a and b**). Interestingly, *fc2 fas2* was tolerant to all tested stresses, making it distinct among all eight mutants.

Thus, there appears to be little cross-tolerance between chloroplast ^1^O_2_ and other abiotic stresses. This conclusion agrees with a transcriptomic data meta-analysis comparing different transcriptional responses to abiotic responses that produce ROS in cells (Rosenwasser et al., 2013). There, it was concluded that chloroplast ^1^O_2_ produces a unique transcriptional signature, suggesting that it may initiate a distinct signal. As such, the lack of multi-stress cross-tolerance observed in most *fc2 fts*/*fas* mutants may be expected, as chloroplast ^1^O_2_ accumulation is unlikely to produce the same transcriptional response as chloroplast or cytosolic H_2_O_2_ produced during the other stresses we tested. Furthermore, this suggests that this screen was able to identify mutant alleles that specifically affect chloroplast ^1^O_2_ signaling and/or tolerance.

### The fc2 fas2 mutant is broadly stress tolerant

Together, these observations show that most *fc2 fas* mutants were specifically altered in their response to chloroplast ^1^O_2_ stress. However, *fc2 fas2* was an intriguing exception. This mutant also exhibited tolerance to EL, MV, heat, freezing, and carbon starvation stresses (**Figs. 8a-e, 9a-c, 10a-d, and 11a-b)**. The reduced accumulation of ^1^O_2_ in *fc2 fas2* suggests it is not a true signaling mutant. However, it accumulates normal levels of chlorophyll, indicating that it likely retains normal tetrapyrrole synthesis and should be able to produce ^1^O_2_ under cycling light conditions (Woodson et al., 2015). Thus, *fas2* may be affecting the quenching of ROS via excess scavengers or by another form of photoprotection.

However, as the *fc2 fas2* mutant exhibits tolerance to non-^1^O_2_-producing stresses, it suggests that the general stress tolerance achieved does not explicitly involve chloroplast ^1^O_2_ or ^1^O_2_-signaling. *fc2 fas2* is a GA-sensitive dwarf mutant (**Fig. 5c),** and a reduction in GA signaling and growth can convey general stress tolerance (Colebrook et al., 2014). However, GA-signaling was pharmacologically uncoupled from the PCD suppression observed in *fc2 fas2* (**Figs. 5b, d, and e),** suggesting that *fc2 fas2* may be using additional mechanisms to increase its stress tolerance.

The tolerance to extreme freezing conditions (−20°C) was striking, particularly as *fc2 fas2* tolerates −20°C for an hour without cold priming (**Figs. 9c and d**). The induction of the cold acclimation and ABA-response genes *COR15a* and *COR27* in *fc2 fas2* may offer a clue (**Figs. 7a and b**). These observations suggest that *fc2 fas2* may be constitutively primed to tolerate freezing stress, possibly by increasing the expression of genes involved in such stresses. It will be interesting to test if such mechanisms can confer general stress tolerance in plants and if ^1^O_2_ stress can offer cross-protection to other types of abiotic stresses.

## Conclusions

Here, we report the initial characterization of eight new gain-of-function dominant *fas* mutants that revealed informative lessons about chloroplast ^1^O_2_ signaling in the *fc2* mutant. First, the ability of *fas* mutations to block ^1^O_2_-initiated PCD is stage-specific and points to the possibility that plants can employ different ^1^O_2_ response strategies, depending on the age of the plant. Second, most *fas* mutations do not lead to broad tolerance to ROS or abiotic stress, indicating that they specifically affect ^1^O_2_ signaling to control PCD and retrograde signaling. This further supports the hypothesis that chloroplast ^1^O_2_ stress induces a unique signal with limited crosstalk with other known stress signaling pathways. Finally, our results implicate SA and JA signaling in the chloroplast ^1^O_2_ response in *fc2* mutants, further connecting these two hormones as important players in chloroplast-mediated PCD and retrograde signaling. Further studies characterizing these mutants will undoubtedly clarify these points and provide a deeper understanding of the molecular mechanisms behind these signals.

## Declarations

### Competing interests

The authors declare that they have no competing interests.

### Funding

The authors acknowledge the Division of Chemical Sciences, Geosciences, and Biosciences, Office of Basic Energy Sciences of the U.S. Department of Energy grant DE-SC0019573 awarded to J.D.W and support from the Center for Research on Programmable Plants and the National Science Foundation grant DBI-2019674. M.D.L was supported by the NIH T32 GM136536 training grant and the UA Richard A. Harvill Graduate Fellowship. The funding bodies played no role in the design of the study and collection, analysis, and interpretation of data and in writing the manuscript.

### Authors’ contributions

MDL and JDW planned and designed the research. MDL performed all screens, identification and characterization of suppressor mutants, management of plant lineages, RT-qPCR, and development of physiological assays and plant stress treatments. JDW conceived the original scope of the project, managed the project, and performed the initial mutagenesis of the plants. MDL and JDW contributed to data analysis and interpretation, wrote the manuscript, reviewed the manuscript, and approved the final version.

## Acknowledgments

This work is dedicated in memory of Susan Rhodes, whose encouragement and advice to M.D.L. throughout his life and academic career helped make this work possible. The authors also wish to thank Marta Kozlowzka and Rebeca Acevedo-Barboza (U of A) for technical support with crosses, Kamran Alamdari, Emma Gevelhoff, and Sophia Daluisio (U of A) for technical assistance with plant maintenance, and Dr. Samantha Orchard (U of A) for the use of lab space.

## Short legends for supporting information

### Supplemental Figures

Figure S1. Baseline chlorophyll fluorescence measurements of plants used in this study

Figure S2. Singlet oxygen accumulation in *fc2 fas* mutants

Figure S3. Experimental design: Activation tagging *fc2* suppressor screen

Figure S4. Adult phenotypes of Class I, Class II, and Class III *fc2* suppressor (*fts*) mutants

Figure S5. Eight *fc2 activation-tagged suppressor* (*fas)* mutants display a dominant cell death suppression phenotype

Figure S6. Analysis of senescence-associated gene expression in *fc2 fas* mutants

Figure S7. Testing the tolerance of *fc2 fas* mutants to heat and freezing stresses

### Supplemental Tables

Table S1. *Arabidopsis thaliana* mutants used in this study

Table S2. Primers used in the study

Table S3. RT-qPCR analysis of ROS-stress response marker genes

Table S4. RT-qPCR analysis of growth hormone response marker genes

Table S5. RT-qPCR analysis of growth hormone response marker genes

Table S6. RT-qPCR analysis of growth hormone response marker genes

Table S7. Classes of *plastid ferrochelatase two suppressor* (*fts*) mutations based on seedling phenotypes

Table S8. Adult phenotypes of select *fc2 fts* mutants in this study

Table S9. Summary of tolerance/sensitivity of *fc2 fts* and *fc2 fas* mutants to stress

## Notes

### Competing Interest Statement

The authors have declared no competing interest.

