## Supplemental information for "A genetic screen for dominant chloroplast reactive oxygen species signaling mutants reveals life stage-specific singlet oxygen signaling networks"

### Supplemental Figures

Figure S1. Baseline chlorophyll fluorescence measurements of plants used in this study

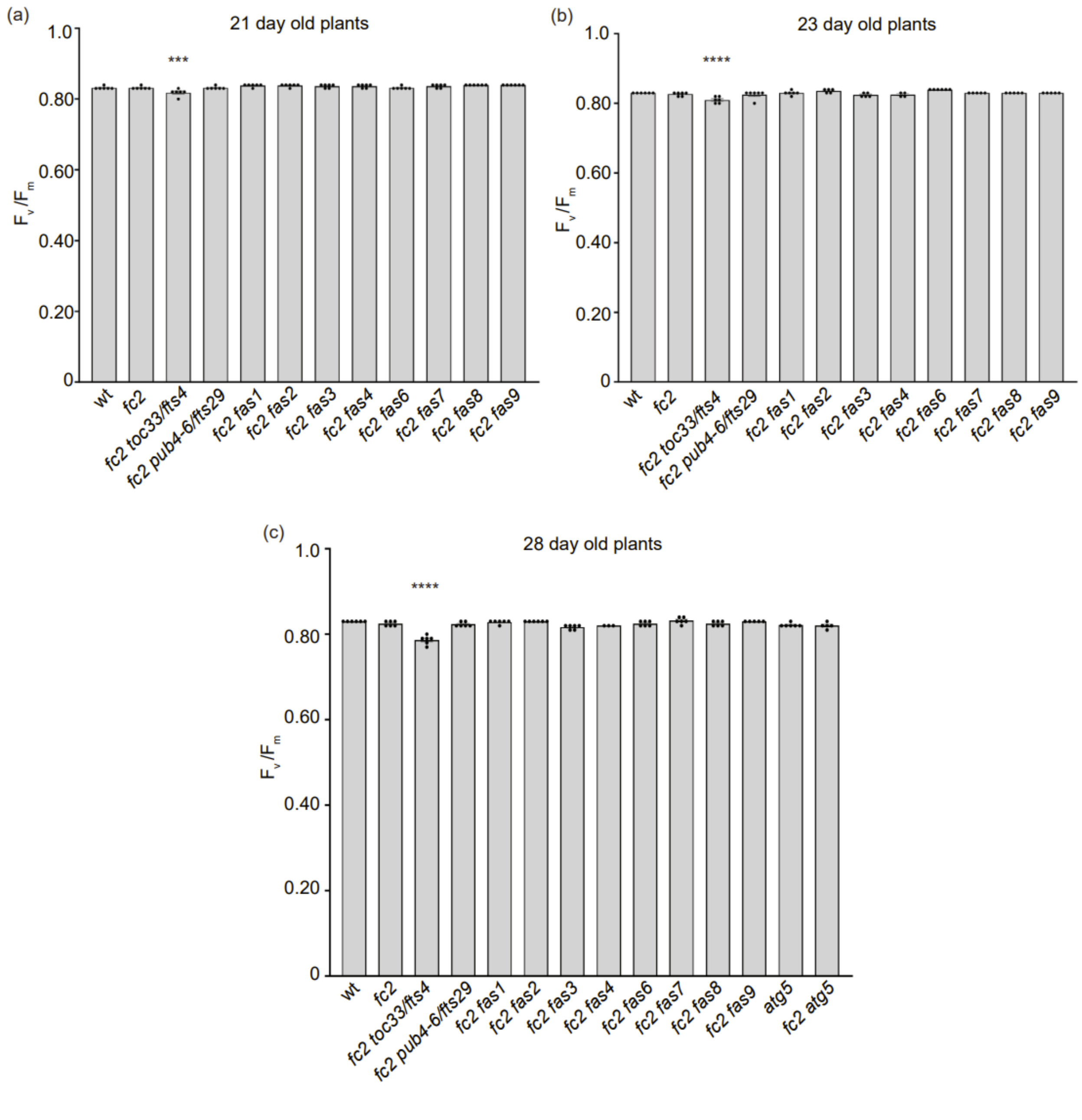

Chlorophyll fluorescence was measured in untreated 21 (a), 23 (b), and 28 (c) day-old plants grown under constant light (24h) conditions to establish a general baseline of untreated and unstressed plants. This is the starting age of plants used in stress treatments in this study. In nearly all cases, plants were within the previously reported $F_v/F_m$ range expected for unstressed plants of ~0.81-0.84 (Bjorkman and Demmig, 1987, Murchie and Lawson, 2013). The only exception was *fc2 toc33*, which consistently had a starting $F_v/F_m$ value of ~0.78-0.81, which was statistically lower than *fc2*. All quantification of $F_v/F_m$ analyses were tested using a one-way ANOVA and Dunnett's multiple comparisons post hoc to compare variation between genotypes relative to *fc2* (*** $P \leq 0.001$, **** $P \leq 0.0001$). n ≥ 6 plants. Error bars = +/- SEM. Closed circles indicate individual data points.

Figure S2. Singlet oxygen accumulation in *fc2 fas* mutants

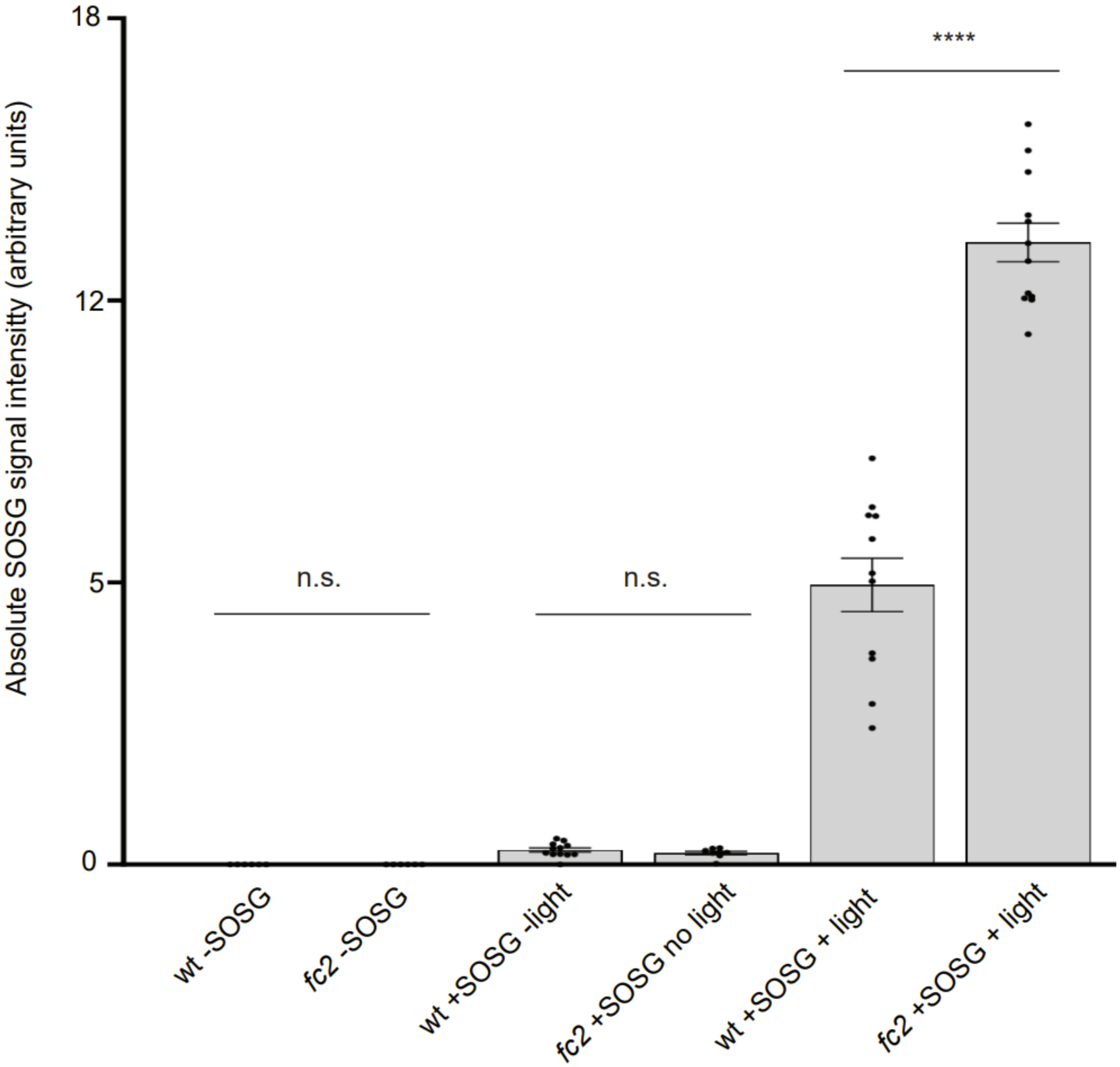

Singlet oxygen ($^1O_2$) accumulation was monitored in 24-day-old leaf tissue using a singlet oxygen sensor green (SOSG). Plants were grown in constant light conditions for 21 days and then transferred to cycling light conditions (16h light/8h dark) for 3 days. Leaf disks were collected and infiltrated with SOSG or a mock control (buffer solution without SOSG). To test the light dependence of $^1O_2$ production in leaf disks and to control for the background fluorescence of SOSG treatment, SOSG-treated leaf disks were either exposed to three hours of white light (120 μmol $m^{-2}$ $sec^{-1}$ white LED light at 21 °C) or kept in dark (at 21 °C) prior to imaging. (a) Quantification of leaf disk SOSG signal. Quantification of SOSG signal was tested with a student's t-tests to compare variation between wt and *fc2 in* each respective group; no SOSG, SOSG -light, SOSG +light (**** = $P \leq 0.0001$). $n \geq 6$ leaf disks from replicate plants. Error bars = +/- SEM. Closed circles indicate individual data points.

Figure S3. Experimental design: Activation tagging *fc2* suppressor screen

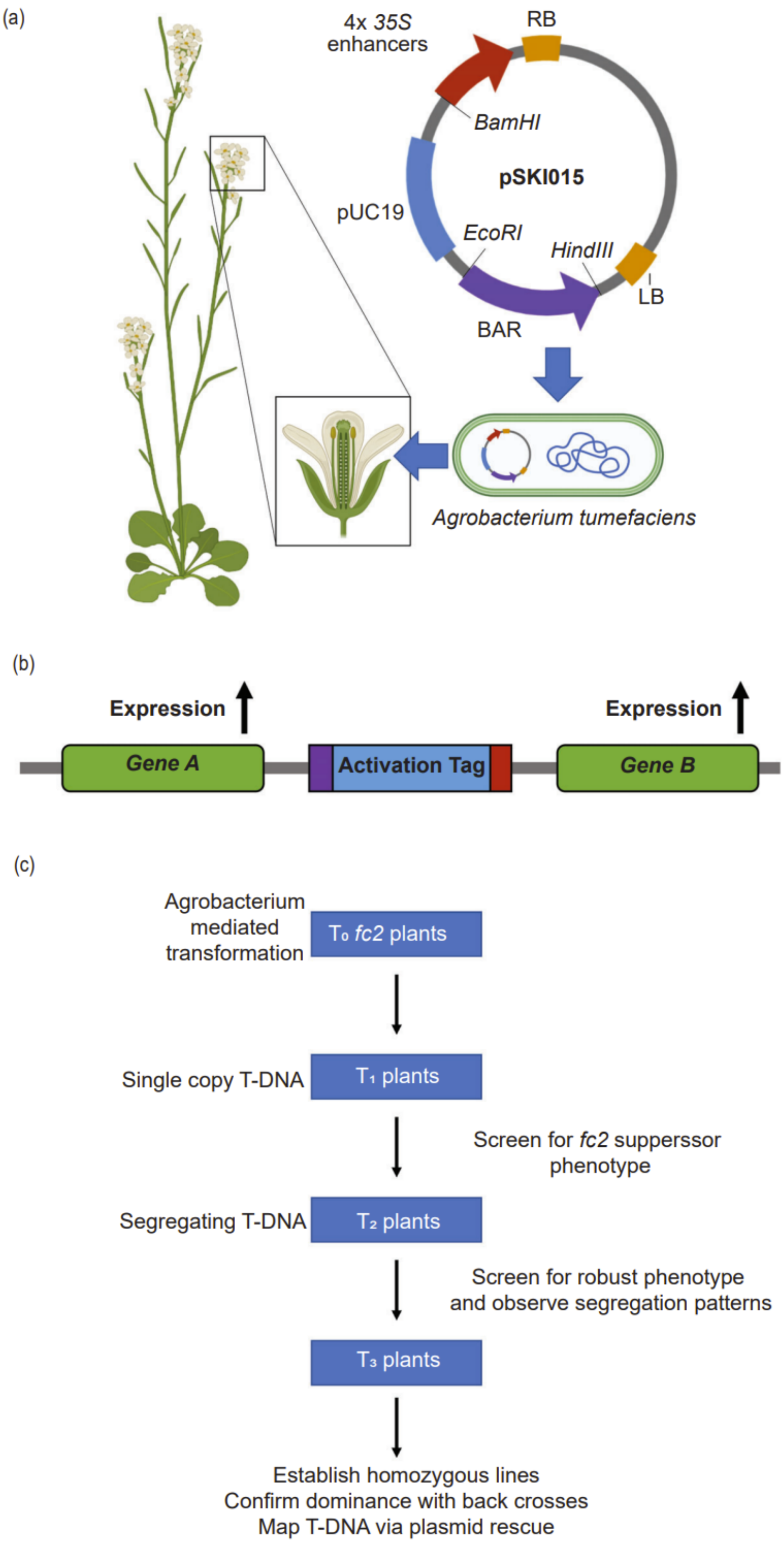

Activation tagging was used to generate dominant *fc2* suppressor mutants. (a) The activation tagging construct pSKI015 (Weigel et al., 2000), which contains four *35S* enhancer sequences, was introduced to the *fc2* mutant background using the *Agrobacterium*-mediated floral dip transformation method. (b) Model depicting the positional effect activation tags have on the expression of adjacent endogenous genes. (c) Workflow of activation tagging screen for dominant *fas* mutants.

Figure S4. Adult phenotypes of Class I, Class II, and Class III *fc2* suppressor (*fts*) mutants

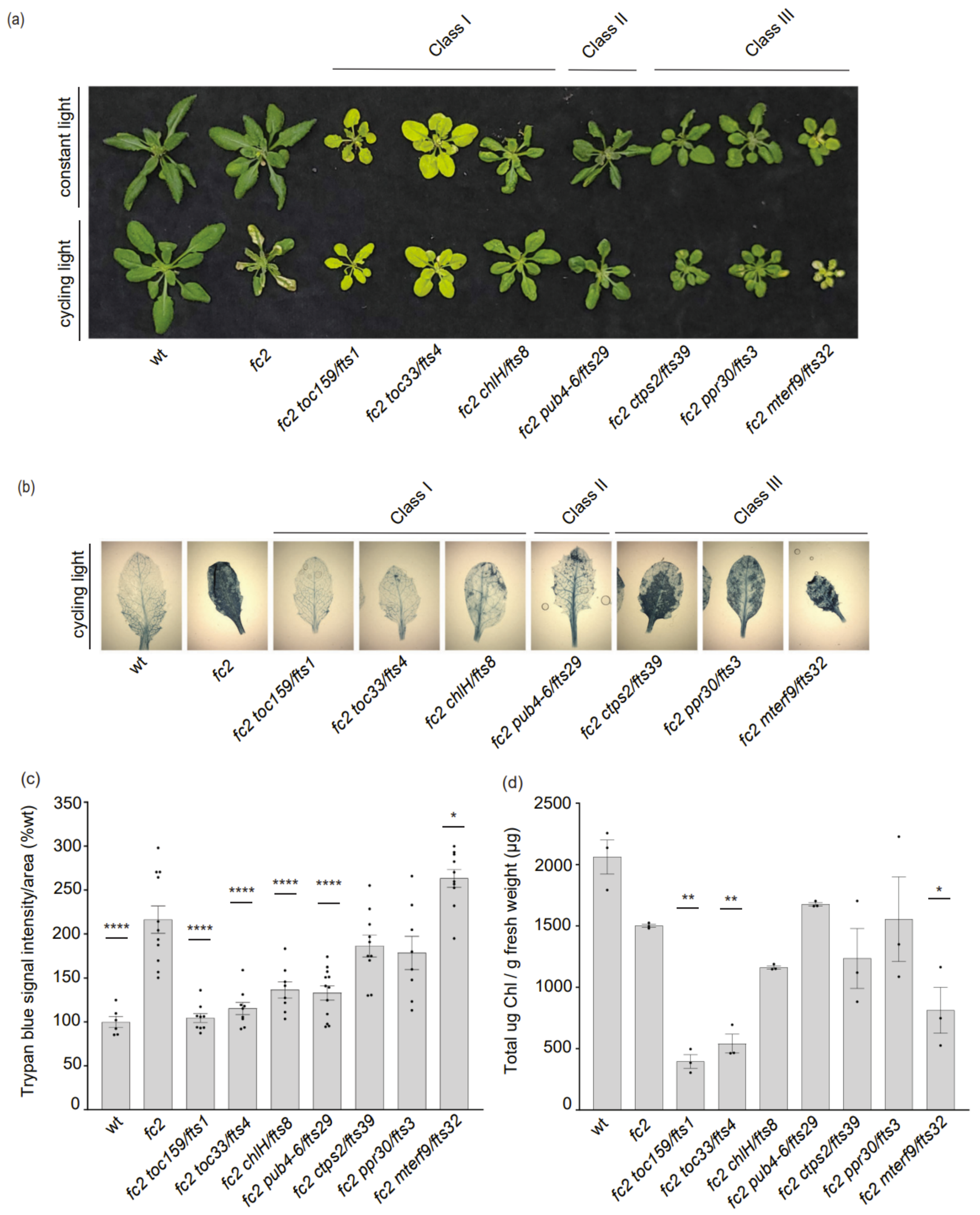

Assessment of adult Class I, Class II, and Class III *fc2 suppressor* (*fts*) mutant phenotypes for the establishment of criteria used for selection of *fas* mutants in this activation tagging screen. Plants

were grown for 21 days in constant light conditions or for 14 days in constant light conditions and then shifted to cycling light conditions (16h light/8h dark) for 7 days. (a) Photograph illustrating the visual phenotypes of representative *fts* mutant plants under both conditions. (b) Representative images of leaves collected from plants grown under cycling light conditions and stained with trypan blue. The dark blue color indicates dead tissue. (c) Quantification of trypan blue stains shown in b ($n \geq 6$ leaves from replicate plants). (d) Total chlorophyll content (µg/g fresh weight) of 21-day-old plants grown in constant light conditions ($n = 3$ biological replicates). Trypan blue stain and chlorophyll content quantification were tested using a one-way ANOVA, and a Dunnett's multiple comparisons post hoc was used to test variation between genotypes relative to *fc2* (* = $P \leq 0.05$, ** = $P \leq 0.01$, **** = $P \leq 0.0001$). Error bars = +/- SEM. Closed circles indicate individual data points.

Figure S5. Eight *fc2 activation-tagged suppressor* (*fas*) mutants display a dominant cell death suppression phenotype.

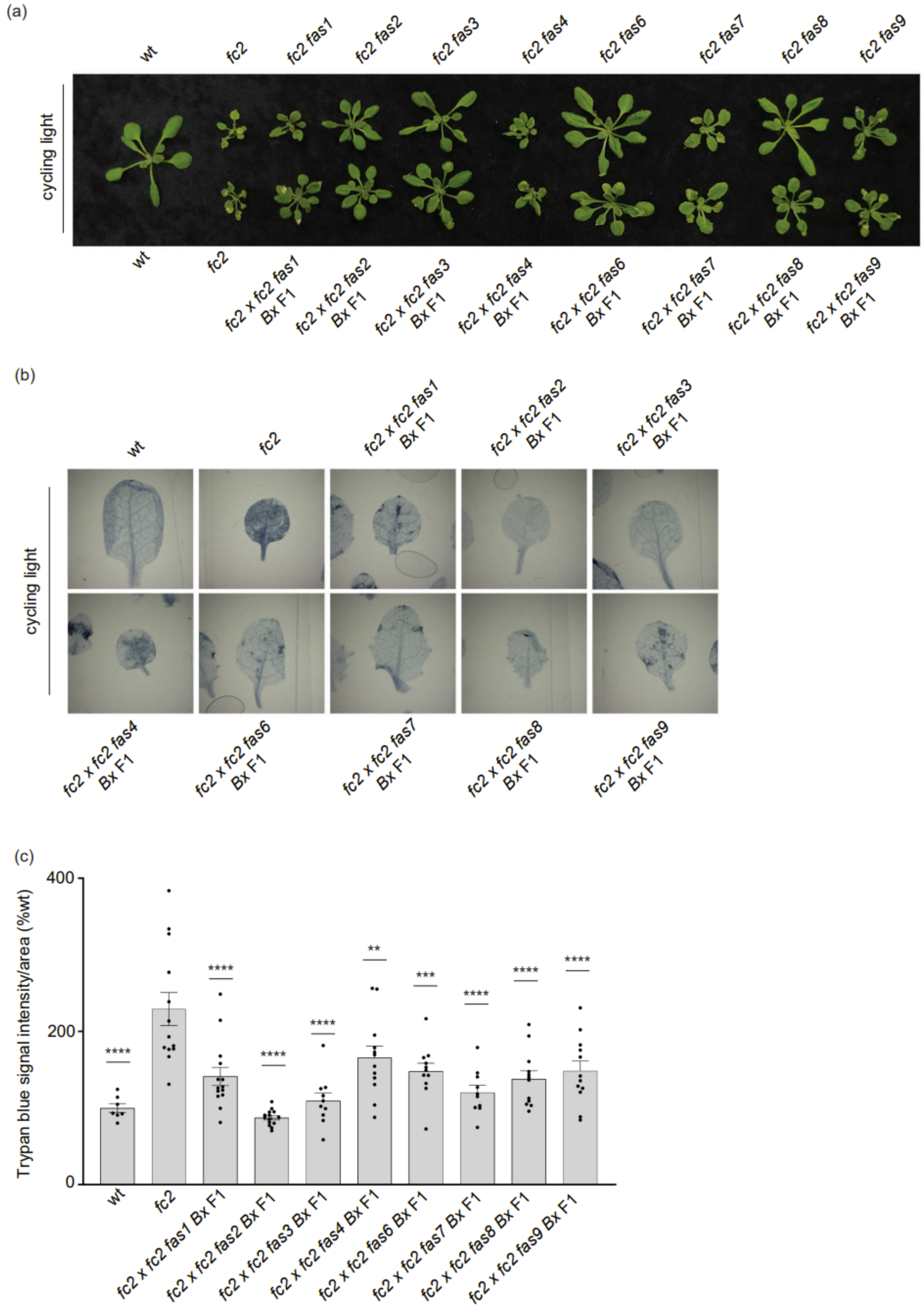

*fas* mutations were assessed for dominance. To this end, homozygous *fc2 fas* mutants (♂) were backcrossed to the parental *fc2* (♀) line. $F_1$ plants were grown for 21 days in constant light conditions or for 14 days in constant light conditions and then transferred to cycling light (16h light/8h dark) conditions for 7 days. (a) Photograph illustrating the visual phenotypes of representative *fc2 fas* mutants and their respective $F_1$ generation backcrosses grown under cycling light conditions. (b) Representative images of leaves collected from plants grown under cycling light conditions and stained with trypan blue. The dark blue color indicates the cell death. (c) Quantification of trypan blue stains shown in b (n ≥ 6 leaves from replicate plants). Trypan blue stain quantification was tested using a one-way ANOVA, and Dunnett's multiple comparisons post hoc was used to test variation between genotypes relative to *fc2* (** = $P \leq 0.01$, *** = $P \leq 0.001$, **** = $P \leq 0.0001$). Error bars = +/- SEM. Closed circles indicate individual data points.

<u>Figure S6. Analysis of senescence-associated gene expression in *fc2 fas* mutants</u>

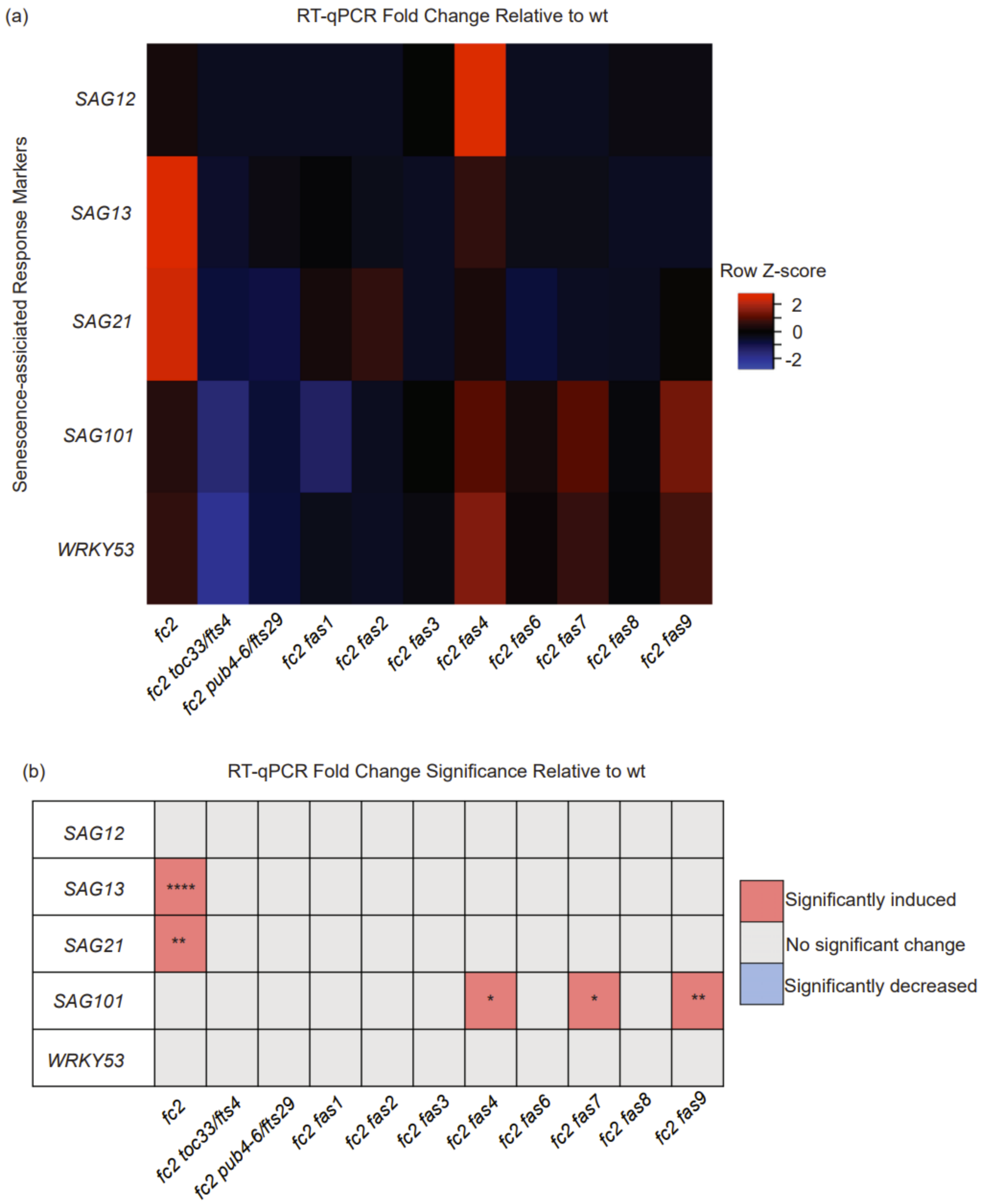

Senescence marker gene expression was measured via RT-qPCR. Transcript abundance of senescence-associated genes was monitored in 21-day-old plants grown for 14 days in constant light conditions and then transferred to cycling light conditions (16h light /8h dark) for seven days. (a) A heatmap table summarizing senescence-associated gene transcript abundance fold change relative to wt. Shades of red indicate increased transcript abundance; shades of blue indicate decreased transcript abundance. (b) Table reporting the significance of difference in senescence marker transcript abundance relative to wt. All quantification of qPCR analyses were tested using a one-way ANOVA and Dunnett's multiple comparisons post hoc to compare variation between genotypes relative to wt (* = $P \leq 0.05$, ** = $P \leq 0.01$, **** = $P \leq 0.0001$). n = 3 biological replicates.

Figure S7. Testing the tolerance of *fc2 fas* mutants to heat and freezing stresses

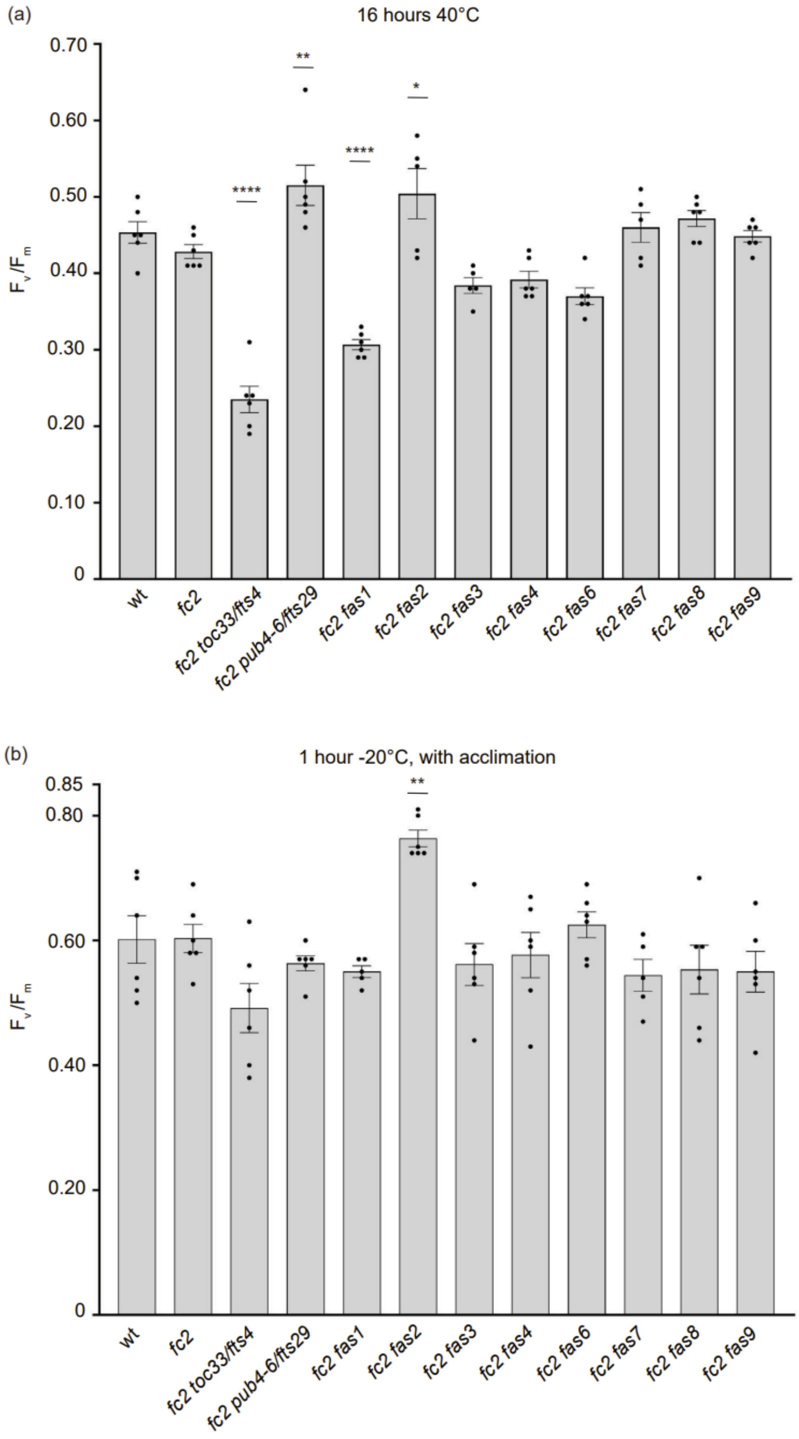

*fc2 fas* mutants were tested for their tolerance to different temperature stresses. Plants were grown for 21 days in constant light conditions and then exposed to heat stress (40°C) for 16h or to freezing stress (-20°C). In the case of freezing stress, plants were first acclimated to cold (16h at 4°C). (a) Quantification of maximum photosynthetic efficiency ($F_v/F_m$) one hour after a 16h 40°C treatment ($n \geq 4$ plants). (b) Quantification of $F_v/F_m$ values taken from cold-acclimated plants, two hours after a 1h -20°C treatment ($n \geq 4$ plants). All $F_v/F_m$ values were tested with a one-way ANOVA and Dunnett's multiple comparisons post hoc to compare variation between genotypes relative to *fc2* (* = $P \leq 0.05$, ** = $P \leq 0.01$, **** = $P \leq 0.0001$). Error bars = +/- SEM. Closed circles indicate individual data points.

**Supplemental Tables**

Table S1: *Arabidopsis thaliana* mutants used in this study

| Mutant | Gene | Locus | Mutation | Change in Protein or Transcript | Citation |
|---|---|---|---|---|---|
| ***fc2-1* (*fc2*)** | *FC2* | *At2G30390* | GABI_766H08 T-DNA in 5'UTR | Reduced accumulation of *FC2* transcript | (Woodson et al., 2011) |
| ***toc159* (*fts1*)** | *TOC159/PPI2* | *At4g02510* | c1109262t | Q1472stop | (Woodson et al., 2015) |
| ***toc33* (*fts4*)** | *TOC33/PPI1* | *At1g02280* | c449839t | Splice change | (Woodson et al., 2015) |
| ***gun5* (*fts8*)** | *GUN5* | *At5g13630* | c4391745t | G113E | (Woodson et al., 2015) |
| ***pub4-6* (*fts29*)** | *PUB4* | *At2g23140* | c9847535t | G255R | (Woodson et al., 2015) |
| ***ctps2-5* (*fts39*)** | *CTPS2* | *At3G12670* | c4023429t | Splice defect | (Alamdari et al., 2021) |
| ***ppr30-1* (*fts3*)** | *PPR30* | *At3g23020* | c8177451t | D765N | (Alamdari et al., 2020) |
| ***mterf9-3* (*fts32*)** | *mTERF9/mTERF31* | *At5g55580* | c22516932t | R368stop | (Alamdari et al., 2020) |
| ***atg5-1*** | *ATG5* | *At5G17290* | SAIL_129_B07 T-DNA intron 4 of 8 | Null | (Thompson et al., 2005) |

Table S2: Primers used in this study

| Gene | Oligo | Sequence |
|---|---|---|
| **Genotyping Primers** | | |
| GABI_766H08 / FC2 - LP | For. JP283 | GAGCAACGCCAAACATAGAAG |
| GABI_766H08 / FC2 - RP | For. JP284 | TCAAAGGCAATGAATGTTTCC |
| GABI-KAT T-DNA RB 03144 | Rev. JP285 | GTGGATTGATGTGATATCTCC |
| pSKI015-790F (T-DNA genotyping probe) | For. WLO1676 | GCGCTCTATCATAGATGTCGCT |
| pSKI015-1974R (T-DNA genotyping probe) | Rev. WLO1677 | TCCGCAGCCATTAACGACTT |
| pSKI015-2261F (T-DNA genotyping probe) | For. WLO1678 | CCCCCGCTGGTATCAAAAGT |
| pSKI015-2972R (T-DNA genotyping probe) | Rev. WLO1679 | ATAGTCCTGTCGGGTTTCGC |
| pSKI015-3772F (T-DNA genotyping probe) | For. WLO1680 | CAGTGCTGCAATGATACCGC |
| pSKI015-4962R (T-DNA genotyping probe) | Rev. WLO1681 | TCAAGCTCTAAATCGGGGGC |
| **RT-qPCR primers** | | |
| **Standard (RT-qPCR)** | | |
| ***AT3G18780 (ACTIN2)*** | For. JP199 | GCACTTGCACCAAGCAGCAT |
| | Rev. JP200 | CCTTTCAGGTGGTGCAACGAC |
| **Stress Response Markers (RT-qPCR)** | | |
| ***fc2*-stress (based on seedlings)** | | |
| ***AT1G52560 (HSP26.5)*** | For. WLO2184 | CGAGCTTATCGTTGCCTGAT |
| | Rev. WLO2185 | CTCCGCCTTAATGTCCTCAA |
| ***AT4G10250 (HSP22)*** | For. JP583 | GTGGCTCTGTCTCCAGCAA |
| | Rev. JP584 | TGCCCTTCTGCTGTTTCTTT |
| ***AT3G56710 (SIB1)*** | For. WLO2182 | CAACCGGAGCCCATCTATT |
| | Rev. WLO2183 | GGAGAAAGGTTGTGGTCGTC |
| **Singlet oxygen($^1O_2$)-stress** | | |
| ***AT3G61190 (BAP1)*** | For. WLO2186 | ATTGATGGATACGGTGGCCG |
| | Rev. WLO2187 | CAGACCCCAAACCGGAACTC |
| ***AT3G28580 (ATPase)*** | For. WLO2188 | GAAGATCGGAAAAGCGTGGAA |
| | Rev. WLO2189 | CCGGGTGGTCCAAACAAAAG |
| ***AT5G64870 (NOD)*** | For. JP340 | GCTGATGCTGCCTTCTATTCAA |
| | Rev. JP341 | TGCGACAAGTCCCTCTGCA |
| **General reactive oxygen species (ROS)-stress** | | |

| | | |
|---|---|---|
| ***AT5G59820 (ZAT12)*** | For. WLO2190 | GCGTTGGTTACACGCGCTT |
| | Rev. WLO2191 | CTTCAACGTAGTCACCGTGGG |
| ***AT4G37370 (CYC81D8)*** | For. WLO2192 | AATGGGCATTGTCGAACGTG |
| | Rev. WLO2193 | TCGCCTTGTTCAATACATCCG |
| ***AT1G17170 (GST)*** | For. JP1126 | GAAGCAGCCAAGGAGTTAATCG |
| | Rev. JP1127 | AAGCTCAGACTCTAGCGTCTTG |
| **Hydrogen ($H_2O_2$) peroxide-stress** | | |
| ***AT5G01600 (FER1)*** | For. JP342 | CACCCAGCTAAGGATGATCGG |
| | Rev. JP343 | TGGTCGAAATGCCAAACTCC |
| ***AT1G07890 (APX1)*** | For. JP1122 | ATTACGCTGAGGCCCACATG |
| | Rev. JP1123 | AAGCATCAGCAAACCCAAGC |
| **Growth hormone response markers (RT-qPCR)** | | |
| **Gibberellin (GA)** | | |
| ***AT1G75750 (GASA1)*** | For. WLO2244 | ACGGAAACTACGACAAGTGC |
| | Rev. WLO2245 | TCTTCTTATGGGCACTTGCG |
| ***AT1G15550 (GA3OX1)*** | For. WLO2246 | ACGCAAGCTTAAGTCTGCTC |
| | Rev. WLO2247 | AACCTTCGGACCACATTTGC |
| **Brassinosteroid (BR)** | | |
| ***AT3G46290 (HERK1)*** | For. WLO2252 | TTCAGCCATTGGTTCGTTGC |
| | Rev. WLO2253 | TCGAAAACGGCATCCAAGTC |
| ***AT1G30570 (HERK2)*** | For. WLO2254 | AAGCTTGGCTTCGCAAACAG |
| | Rev. WLO2255 | AAGAGCAGATCCTGTTTGGC |
| **Auxin (IAA)** | | |
| ***AT2G38120 (AUX1)*** | For. WLO2240 | ATGCTTTCGTGGTGGTTTGG |
| | Rev. WLO2241 | GCTGCAGCTGGTTTACATTG |
| ***AT1G15580 (IAA5)*** | For. WLO2242 | TGCTTCCGCTCTGCAAATTC |
| | Rev. WLO2243 | AACATCTCCAGCAAGCATCC |
| **Cytokinin (CK)** | | |
| ***AT1G10470 (ARR4)*** | For. WLO2248 | AATTGAATCTGCGCCGTTGG |
| | Rev. WLO2249 | AAATTCCAGAGCACGCCATC |
| ***AT3G48100 (ARR5)*** | For. WLO2250 | TCTACTCGCAGCTAAAACGC |
| | Rev. WLO2251 | TCAGGACATGCATGTGTGTG |
| **Stress hormone response markers (RT-qPCR)** | | |
| **Salicylic acid (SA)** | | |
| ***AT2G14610 (PR1)*** | For. WLO1771 | GTGCTCTTGTTCTTCCCTCGA |
| | Rev. WLO1772 | CCCACGAGGATCATAGTTGCA |
| ***AT3G57260 (PR2)*** | For. WLO2226 | GATCGTTGGAAATCGTGGTG |
| | Rev. WLO2227 | TAGCTTTCCCTGGCCTTCTC |

| | | |
|---|---|---|
| ***AT1G75040 (PR5)*** | For. WLO2258 | TTGCAAATACGCAGGCTGTG |
| | Rev. WLO2259 | ACGGCAGCAATATTGATCCG |
| **Jasmonic acid (JA)** | | |
| ***AT3G12500 (PR3)*** | For. WLO2150 | TGCAACTGTCGTGGAACTAC |
| | Rev. WLO2151 | AGAACCAAATCGCGGCTTTG |
| ***AT3G04720 (PR4)*** | For. WLO2256 | AGTGCTTATTGCTCCACGTG |
| | Rev. WLO2257 | AAACACTTGCCGCAAGAAGC |
| ***AT1G19180 (JAZ1)*** | For. WLO2264 | TGCAACCAAACCAACCAACC |
| | Rev. WLO2265 | TGTTGCTGTTGCTTCTCTGC |
| **Abscisic acid (ABA)** | | |
| ***AT2G42540 (COR15a)*** | For. WLO1887 | AACGAGGCCACAAAGAAAGC |
| | Rev. WLO1888 | TTCGCTTTCTCACCATCTGC |
| ***AT5G42900 (COR27)*** | For. WLO2152 | AAAGGCGAAAACGGAAGCTC |
| | Rev. WLO2153 | TGAGTGGAACAACCTGATCGG |
| ***AT5G06760 (LEA4-5)*** | For. WLO1775 | CTTGGAGGAAAAGGCGGAGA |
| | Rev. WLO1776 | ATCCAGTATATCCCCCGCCG |
| **Ethylene (ET)** | | |
| ***AT3G23240 (ERF1)*** | For. WLO1773 | CCATTCTCCGGCTTCTCACC |
| | Rev. WLO1774 | TTCACGGAGCGGTGATCAAA |
| ***AT5G25350 (EBF2)*** | For. WLO2154 | AGCTGTTGGAAATGGTTGCC |
| | Rev. WLO2155 | ATGCAGATTTGGCCAAAGCG |
| ***AT1G01480 (ACS2)*** | For. WLO2262 | TGCTTGCTTCGATGTTGTCC |
| | Rev. WLO2263 | ACCAGCGTTGCTTGTCAAAC |
| **Senescence associated markers (RT-qPCR)** | | |
| ***AT5G45890 (SAG12)*** | For. WLO2269 | GGAGGTGGTTTTGATTTCCA |
| | Rev. WLO2270 | GACATCAATCCCACACAAACA |
| ***AT2G29350 (SAG13)*** | For. WLO1834 | AGCCATGTTGGGAGCAAAAG |
| | Rev. WLO1835 | TGCTTGCCATTCACGTAAGC |
| ***AT4G02380 (SAG21)*** | For. WLO2271 | TGCTTTCGTCTCTCGTGAAC |
| | Rev. WLO2272 | TTCATCACAGCCGAAGCAAC |
| ***AT5G14930 (SAG101)*** | For. WLO1836 | TTGCTCCAAACGCTCAAGTC |
| | Rev. WLO1837 | AACGTCGGTTCGATGGTTTC |
| ***AT4G23810 (WRKY53)*** | For. WLO2273 | GCCGCCGTTTTACAATTACG |
| | Rev. WLO2274 | AAAACGCGGGGAAAGTTGTG |

Table S7: Classes of *plastid ferrochelatase two suppressor* (*fts*) mutations based on seedling phenotypes

| **Suppressor class** | **Block retrograde signal (cycling light conditions)** | **Block cell death? (cycling light conditions)** | **Reduced chlorophyll? (constant light conditions)** | ***fc2 suppressor* (*fts*) mutants** | **Proposed mechanism** |
|---|---|---|---|---|---|
| Class I | Yes | Yes | Yes | *fc2 toc159/ppi2/fts1 (Woodson et al., 2015)*<br>*fc2 toc33/ppi1/fts4 (Woodson et al., 2015, Lemke et al., 2021)*<br>*fc2 toc33/ppi1/fts37 (Woodson et al., 2015)*<br>*fc2 gun5/fts8 (Woodson et al., 2015)*<br>*fc2 gun5/fts16 (Woodson et al., 2015)*<br>*fc2 gun5/fts24 (Woodson et al., 2015)*<br>*fc2 gun5/fts30 (Woodson et al., 2015)* | Decreased tetrapyrrole biosynthesis |
| Class II | Yes | Yes | No | *fc2 pub4-6/fts29 (Woodson et al., 2015, Lemke et al., 2021, Tano et al., 2022)* | Altered protein ubiquitination |
| Class III | Yes | Yes | Yes | *fc2 ctps2-5/fts39 (Alamdari et al., 2021)*<br>*fc2 ppr30-1/fts3 (Alamdari et al., 2020, Tano et al., 2022)*<br>*fc2 ppr30-2/fts38 (Alamdari et al., 2020)* | Impaired plastid gene expression |

| | | | | |
|---|---|---|---|---|
| | | | | *fc2 mterf9-3/fts32 (Alamdari et al., 2020)* |

Seedling phenotypes caused by the *plastid ferrochelatase 2 suppressor* (*fts*) mutations (blocked retrograde signaling, blocked programmed cell death (PCD), and reduced total chlorophyll content) led to their placement in three classes. Class I suppressor mutations block conditional PCD in cycling light conditions and generally lead to reduced total chlorophyll content in constant light conditions. These mutations directly or indirectly affect tetrapyrrole biosynthesis and accumulation, thereby reducing singlet oxygen ($^1O_2$) production. Class II suppressor mutations block conditional PCD in cycling light conditions, but do not reduce total chlorophyll content in constant light conditions. This class presumably acts by a mechanism downstream of chloroplast $^1O_2$ accumulation and can involve the cytoplasmic ubiquitination machinery. Class III suppressor mutations generally block conditional PCD in cycling light conditions and generally reduce total chlorophyll content in constant light conditions. This class of suppression is achieved via impairments to plastid gene expression that is hypothesized to reduce expression of a plastid-encoded signaling factor. This also results in delayed chloroplast development. All mutant classes also reduce chloroplast retrograde signaling suggesting there is a strong link between this signaling and PCD.

Table S8: Adult phenotypes of select *fc2 fts* mutants in this study

| Suppressor class | Block cell death? (cycling light conditions) | Reduced chlorophyll? (constant light conditions) | *fc2 suppressor* (*fts*) mutants tested |
|---|---|---|---|
| Class I | Yes | Yes | *fc2 toc159/fts1 (Woodson et al., 2015)*<br>*fc2 toc33/fts4 (Woodson et al., 2015)*<br>*fc2 chlH/fts8 (Woodson et al., 2015)* |
| Class II | Yes | No | *fc2 pub4-6/fts29 (Woodson et al., 2015)* |
| Class III | No | No | *fc2 ctps2-5/fts39 (Alamdari et al., 2021)*<br>*fc2 ppr30-1/fts3 (Alamdari et al., 2020)*<br>*fc2 mterf9-3/fts32 (Alamdari et al., 2020)* |

In this study, we characterized the adult phenotypes of previously published *fc2 fts* mutants that had only been characterized in the seedling stage. We observed that Class I suppressor mutations block PCD in cycling light conditions and generally reduce total chlorophyll content in constant light conditions. Class II suppressor mutations block conditional PCD in cycling light conditions, but do not reduce total chlorophyll content in constant light conditions. Class III suppressor mutations do not block conditional PCD to a significant degree in cycling light conditions, and do not generally reduce total chlorophyll content in constant light conditions. It should be noted that while we report here that Class III suppressor mutations fail to significantly block PCD under stringent conditions in the adult stage (~120 µmol photons $m^{-2}$ $sec^{-1}$, LED light), suppression of PCD has been observed under more permissive conditions (~100 photons µmol $m^{-2}$ $sec^{-1}$, fluorescent light), as has been previously reported with *fc2 ppr30-1/fts3* (Tano et al., 2022). All adult *fc2 fts* data reported in this work are in **Figs. S4a-d**.

Table S9: Summary of tolerance/sensitivity of *fc2 fts* and *fc2 fas* mutants to stress

| Stress | Treatment | *fc2 toc33* | *fc2 pub4-6* | *fc2 fas1* | *fc2 fas2* | *fc2 fas3* | *fc2 fas4* | *fc2 fas6* | *fc2 fas7* | *fc2 fas8* | *fc2 fas9* |
|---|---|---|---|---|---|---|---|---|---|---|---|
| **cycling light** | 7 days (Trypan blue) | T | T | T | T | T | T | T | T | T | T |
| **excess Light (1450-1500 µmol $m^{-2}$ $sec^{-1}$)** | 24h (Lesions) | S | T | T | T | - | - | - | T | T | - |
| | 24h (Trypan blue) | - | T | T | T | - | T | - | T | T | T |
| | 24h ($F_v/F_m$) | S | T | T | T | - | S | - | - | T | - |
| **methyl viologen ($O_2^-$, $H_2O_2$)** | 20 µM (Lesions) | - | - | - | T | - | - | - | - | - | - |
| | 20 µM ($F_v/F_m$) | T | - | - | T | - | - | - | - | - | - |
| **heat (40° C)** | 16h ($F_v/F_m$) | S | T | S | T | - | - | - | - | - | - |
| | 24h ($F_v/F_m$) | S | T | - | T | - | - | - | - | - | - |
| **freeze (-20° C)** | 1h +acclimation ($F_v/F_m$) | - | - | - | T | - | - | - | - | - | - |
| | 1h -acclimation ($F_v/F_m$) | S | - | - | T | - | - | - | S | S | - |
| **carbon starvation (dark induced)** | 5 days ($F_v/F_m$) | T | - | - | T | - | - | - | T | - | - |
| | 7 days ($F_v/F_m$) | - | - | - | - | - | - | - | T | - | - |

Table summarizing the tolerance or sensitivity of *fc2 fts* and *fc2 fas* mutants to a wide range of stresses. An orange hue and "T" indicate tolerance, a blue hue and "S" indicates sensitivity, and a grey hue and "–" indicate no significant difference, all relative to *fc2*. The *fc2* mutant is sensitive to cycling light stress but showed no pattern of tolerance or sensitivity to any of the additional stresses tested, relative to wt. Trypan blue = cell death stain, lesions = lesion count, $F_v/F_m$ = maximum quantum efficiency of photosystem II. See main text for details on how tolerance and sensitivity were assessed.

### Gene Expression Data Tables

All RT-qPCR data used to generate heatmaps (Tables S3-6) can be found in the associated Microsoft Excel file.

### Supplemental Methods

#### <u>Standard growth conditions</u>

Seedlings (Woodson et al., 2015) and adult plants (Lemke et al., 2021) were germinated and grown as previously described. Briefly, seeds used in growth experiments were sterilized overnight using chloride gas (generated by mixing concentrated HCl (4 ml) and bleach (150 ml) in a domed desiccator) and then resuspended in a 0.1% agar solution. Resuspended seeds were then spread on plates containing 1x Linsmaier and Skoog medium pH 5.7 (Caisson Laboratories North Logan, UT) in 0.6% micropropagation type-1 agar powder (PlantMedia, CAS:9002-18-0). Plates were then stratified in the dark at 4°C for three to five days and then germinated and grown in constant light (24h light) conditions (or diurnal cycling light (6h light/18h dark) conditions in the case of seedling stress tests) in fluorescent light chambers (Percival® model CU-36L5 plant tissue culture chamber) with a light intensity of ~100-115 photons μmol photons $m^{-2}$ $sec^{-1}$ at 21°C. For adult plant experiments, seven-day-old seedlings grown in constant light conditions were carefully transferred to soil (PRO-MIX LP15) supplemented with fertilizer (Jack's Classic All Purpose fertilizer, 6.7 mL of 300 g/L solution per flat), and growth was continued under similar conditions as seedlings (~110-120 μmol photons $m^{-2}$ $sec^{-1}$ at 21°C) in a reach-in LED plant growth chamber (Hettich PRC 1700), set to constant light (24h light) or diurnal cycling light (16h light/8h dark) conditions. Photosynthetically active radiation was measured using a LI-250A light meter with a LI-190R-BNC-2 Quantum Sensor (LiCOR).

Confirmation of T-DNA mutant lines by PCR genotyping

All PCR was performed as previously described (Lemke et al., 2021) using Promega GoTaq® Master Mix, with an initial denaturation step at 93°C for 3 min, followed by a 30s denaturation step at 93°C, a 30s annealing step at 60°C, and a 2 min elongation step at 72°C, cycled 35 times, and then concluded with a final elongation step at 72°C for 7 min. These thermocycler settings were used for all primer sets. This resulted in DNA fragments between ~0.5-1.5 kb in length, which were separated on a 1% agarose gel for imaging. Primers used for PCR genotyping are listed in **Table S2**.

Biomass measurements

Biomass was assessed as previously described (Lemke et al., 2021). Here, green shoot tissue (above ground) was collected for biomass measurements. Replicates of each line and growth condition were stored in envelopes for desiccation. Plant tissue was desiccated in a 65°C desiccation oven for 48 hours and then weighed.

*Chlorophyll fluorescence measurements*

Chlorophyll fluorescence measurements were conducted as previously described (Lemke et al., 2021) and adapted for adult plants. Briefly, maximum quantum yield of PSII ($F_v/F_m$) measurements were obtained from adult plants grown for at least 21 days under constant light conditions prior to treatments and after exposure to EL, MV, Heat, Freezing, or dark-induced carbon starvation stresses. Plants were dark acclimated in a FluorCam chamber (Closed FluorCam FC 800-C/1010-S, Photon Systems Instruments) for at least 15 minutes. Measurements were taken following the manufacturer's manual. $F_v/F_m$ measurements were taken from each replicate plant consecutively from 4 to 6 replicates in the same pot of soil in the same flat of pots as all other genotypes.

*Chlorophyll measurements*

Plant chlorophyll content was measured as previously described in (Woodson et al., 2015). Here, above-ground shoot tissue from 21-day-old plants was weighed and flash-frozen using liquid nitrogen. Tissue was then disrupted and resuspended in ice-cold 80% acetone via vortexing and incubated in the dark at -20°C for 20 minutes. Samples were then centrifuged (21,130 x *g*) at 4°C for 10 minutes, and the supernatant was transferred to fresh tubes. The remaining pellet was resuspended a second time in ice-cold 80% acetone, and the supernatant was collected as described above. The extracted chlorophyll solution was diluted (1:4) in 80% acetone and then transferred to a cuvette, and the absorbance at A645 and A663 was measured using a spectrometer. Total chlorophyll content was calculated using Arnon's equations (Arnon, 1949).

*Cell death measurements*

Cell death was assessed using trypan blue staining as previously described (Alamdari et al., 2020, Lemke et al., 2021). Here, seedings or adult leaves were collected and transferred to a solution of Trypan Blue (stock solution (1.25 ml phenol, 1.25 ml glycerol, 1.25 ml lactic acid, 0.625 ml $H_2O$, and 0.625 ml trypan blue solution (0.4% w/v) (Corning)) diluted with ethanol 1:2

(v/v)). The solution was then boiled for 1 minute, incubated overnight, and then destained twice with a solution of saturated chloral hydrate (2.5 g per 1 ml water). Chloral hydrate was then removed, and seedlings or adult leaves were stored in a 30% glycerol solution prior to imaging on a dissection scope. Replicates for seedlings were grown on the same plate in the same conditions. For adult plants, true leaves (#'s 3-6) were selected from separate plants. For each replicate, staining intensity was measured across one entire cotyledon (seedlings) or leaf (adult plants) using ImageJ software.

*RNA extraction and RT-qPCR*

Transcript levels were monitored as previously described (Alamdari et al., 2020). Total RNA was extracted from whole seedlings using the RNeasy Plant Mini Kit (Qiagen) and cDNA was synthesized using the Maxima first-strand cDNA synthesis kit for RT-qPCR with DNase (Thermo Scientific) following the manufacturer's instructions. Real-time PCR was performed using the iTAQ Universal SYBR Green Supermix (BioRad) with the SYBR Green fluorophore and a CFX Connect Real-Time PCR Detection System (BioRad). The following 2-step thermal profile was used in all RT-qPCR: 95 °C for 3 min, 40 cycles of 95 °C for 10 s, and 60 °C for 30 s (as per the manufacturer's instructions). All expression data was normalized according to *ACTIN2* expression. Primers used for RT-qPCR are listed in **Table S2**.
